# All-Atom Protein Sequence Design using Discrete Diffusion Models

**DOI:** 10.1101/2025.06.13.659451

**Authors:** Amelia Villegas-Morcillo, Gijs J. Admiraal, Marcel J.T. Reinders, Jana M. Weber

**Affiliations:** Department of Intelligent Systems, Delft University of Technology, Delft, The Netherlands

**Keywords:** Protein Sequence Design, All-Atom Representation, Discrete Diffusion Models, Generative Modeling

## Abstract

Advancing protein design is crucial for breakthroughs in medicine and biotechnology. Traditional approaches for protein sequence representation often rely solely on the 20 canonical amino acids, limiting the representation of non-canonical amino acids and residues that undergo post-translational modifications. This work explores discrete diffusion models for generating novel protein sequences using the all-atom chemical representation SELFIES. By encoding the atomic composition of each amino acid in the protein, this approach expands the design possibilities beyond standard sequence representations. Using a modified ByteNet architecture within the discrete diffusion D3PM framework, we evaluate the impact of this all-atom representation on protein quality, diversity, and novelty, compared to conventional amino acid-based models. To this end, we develop a comprehensive assessment pipeline to determine whether generated SELFIES sequences translate into valid proteins containing both canonical and non-canonical amino acids. Additionally, we examine the influence of two noise schedules within the diffusion process—uniform (random replacement of tokens) and absorbing (progressive masking)—on generation performance. While models trained on the all-atom representation struggle to consistently generate fully valid proteins, the successfully generated proteins show improved novelty and diversity compared to their amino acid-based model counterparts. Furthermore, the all-atom representation achieves structural foldability results comparable to those of amino acid-based models. Lastly, our results highlight the absorbing noise schedule as the most effective for both representations. Data and code are available at https://github.com/Intelligent-molecular-systems/All-Atom-Protein-Sequence-Generation.

**Scientific Contribution:** This work introduces a discrete diffusion-based framework for protein sequence generation using an all-atom representation, enabling the incorporation of non-canonical amino acids and post-translational modifications. Additionally, it provides a comprehensive evaluation pipeline to assess the validity of generated proteins, demonstrating how noise schedules within the diffusion process impact sequence novelty, diversity, and structural foldability.

## 1 Introduction

The ability to successfully design proteins enables transformative solutions for medicine, industry, and environmental sciences [1]. By creating novel proteins or optimizing existing ones, we can develop targeted therapeutics, efficient vaccines, and specialized enzymes for industrial and environmental applications [2]. Proteins are macromolecules composed of long chains of amino acids linked by peptide bonds. The specific sequence of these amino acids dictates how a protein folds into its unique three-dimensional structure, which ultimately determines its function [3]. Protein design involves manipulating either the amino acid sequence [4], the three-dimensional structure [5], or both to achieve desired functional properties [6].

Recent computational advancements, particularly AlphaFold2 [7], have addressed a critical limitation in protein design—structural data scarcity—by enabling accurate structure predictions directly from amino acid sequences [8]. This breakthrough facilitates computational protein design based on sequence information alone. Since experimental techniques for structure determination are resource-intensive and often capture only static snapshots of inherently dynamic proteins [9, 10], structure prediction has become an essential tool for advancing protein design.

Generative models—especially diffusion models—have revolutionized protein design [11, 12]. Diffusion models use a noising process to progressively transform data into noise, and then learn to reverse this process to generate new samples [13–15]. Their success in fields such as computer vision [16] and protein design [5] stems from their ability to produce diverse outputs that can be guided toward specific design objectives. For categorical data like amino acid sequences, discrete denoising diffusion probabilistic models (D3PMs) [17, 18] offer an effective alternative, applying noise through transition matrices characterized by uniform transitions or masking strategies. Protein sequence and small molecule design have been explored using various discrete diffusion models. One method is EvoDiff [19], which employs D3PMs to generate amino acid sequences through a masking-based noising process. Similar approaches, such as discrete flow models (DFMs) [20] and diffusion optimized sampling (NOS) [21], also explore discrete diffusion for canonical amino acid representations. In contrast, the functional-group-based diffusion framework (D3FG) [22] combines discrete and continuous diffusion for small molecule generation conditioned on a protein target, focusing on 25 functional groups—key chemical substructures that drive molecular properties—and single-atom linkers.

Existing protein sequence design methods rely on conventional amino acid-level representations, where proteins are encoded as strings of the 20 canonical amino acids found in nature [4]. This approach reflects biological processes in which ribosomes translate mRNA sequences into polypeptide chains [23]. However, this traditional representation has limitations, such as its inability to account for all possible noncanonical amino acids [24] or residues that undergo post-translational modifications (PTMs) [25], which can diversify protein functions.

To address these limitations, we propose leveraging all-atom chemical representations for protein sequence design. Beyond including non-canonical amino acids, the all-atom sequence representation allows for modeling protein-small molecule interactions within a shared representation space. This capability is particularly valuable for designing protein binding pockets [26] conditioned on interacting small molecules, with applications in antibody engineering, enzyme optimization, and biosensor development [27–29]. Moreover, this approach facilitates the design of optimized ligands targeting specific proteins, advancing applications in protein-conditioned drug discovery [30, 31]. One promising all-atom approach is self-referencing embedded strings (SELFIES) [32], a string-based molecular representation designed for use in generative models. SELFIES can represent molecules of varying complexity, including longer and more intricate structures [33], while ensuring that every generated sequence corresponds to a chemically valid molecule—an essential feature for protein design. All-atom representations have already demonstrated success in generating small molecules [34], polymers [35], and therapeutic peptides [36]. However, for large protein design tasks, only one existing study has employed SELFIES in generative pre-trained transformers (GPTs), training on canonical amino acids and datasets incorporating molecular fragments for non-canonical amino acids [37]. While this method showed promise in generating diverse proteins, it relied on an auto-regressive model that generates sequences sequentially, therefore not benefiting from the efficiency of parallel generation in diffusion models.

This work introduces the first all-atom diffusion model for large protein design, integrating the SELFIES representation within the discrete diffusion D3PM framework. By representing every atom in the protein backbone and side chains, this approach provides greater flexibility for incorporating non-canonical amino acids and PTMs. Additionally, we investigate the impact of different D3PM noising processes, such as uniform and absorbing noise, on protein sequence generation.

A key contribution of this work is the evaluation of the generated all-atom sequences. Although the SELFIES-generated sequences are chemically valid, this alone does not ensure that the resulting molecule is a protein. Therefore, we assess whether the SELFIES strings can be converted into valid proteins, and categorize their amino acids as canonical or non-canonical. This evaluation, combined with analyses of protein sequence novelty, diversity, and structural foldability, offers valuable insights into the functional potential of all-atom-generated proteins. Together, these contributions pave the way for the application and understanding of all-atom representations in discrete diffusion models for protein design.

## 2 Methodology

### 2.1 Dataset

Our diffusion models are trained on the UniRef50 dataset [38], which groups protein sequences with a 50% sequence identity threshold. UniRef50 was selected for its balance between comprehensive coverage and sequence diversity. This ensures the model is exposed to a wide range of protein sequences while avoiding high similarity. We obtained protein sequences from the UniRef50 dataset on May 7th 2024. We kept sequences containing solely the 20 canonical amino acids, since non-standard amino acids are denoted as unknown (X), and can thus not be converted to an all-atom representation. While this decision may bias generation toward canonical amino acids, we find that the all-atom models can, in fact, expand beyond this set and generate sequences containing rare, non-canonical residues. Then, we limited the sequence lengths from 30 to 100 amino acids to avoid excessive sequence length expansion with the all-atom representation. After filtering, the final dataset contains around 14 million protein sequences. We split this dataset into training and validation sets (90*/*10 ratio) with a balanced distribution of protein sequence lengths. In Supplementary Material B, we present our dataset statistics, including the distributions of amino acid and SELFIES sequence lengths, as well as the frequency of amino acid tokens.

### 2.2 Protein Sequence Representations

The input to our generative model is either the amino acid sequence tokenized using the 20 canonical amino acids (single-letter codes), or its all-atom representation given by SELFIES [32] (21 tokens). In our case, SELFIES is preferable over other representations, such as SMILES (simplified molecular-input line-entry system) [39], because of its ability to always generate valid molecules (as detailed in Supplementary Material A.1). Fig. 1 illustrates an example of both all-atom representations for the amino acid Proline, showing how random mutations in the symbols affect the resulting molecule. By using SELFIES, we can process the outputs directly, instead of first validating them chemically. Each amino acid sequence is translated to its corresponding SELFIES representation using the Python packages RDKit [40] and SELFIES [41]. The SELFIES grammar compared to the amino acid tokens is shown in Table 1.

**Fig. 1:**
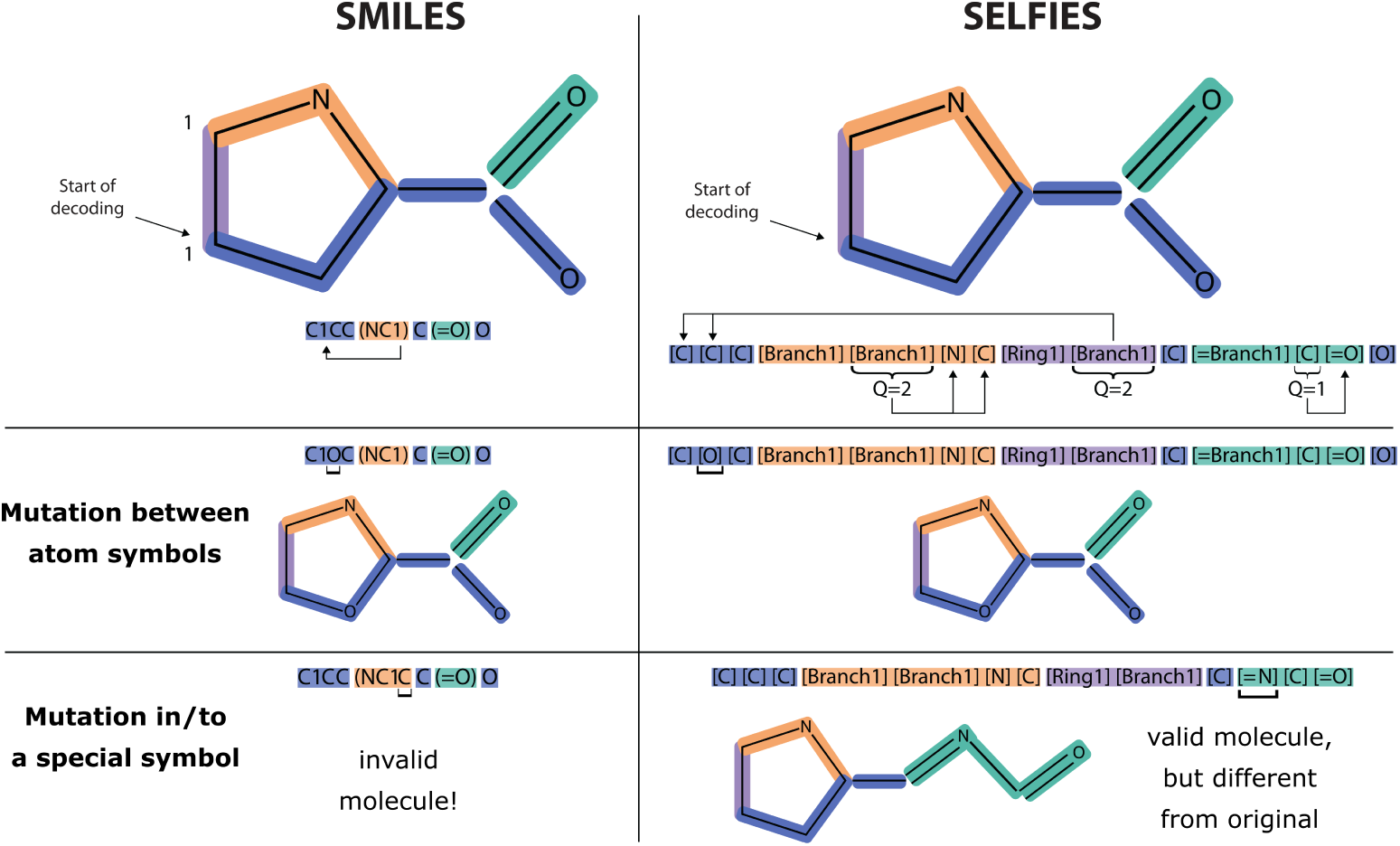
Comparison of SMILES and SELFIES representations for the amino acid Proline. Both examples show how branching and ring formation are handled in their respective representations. In SELFIES, overloaded symbols are annotated with their length or connectivity (Q) to handle these structural features. Decoding starts at the left in both representations and leads to the same canonical SMILES string, which we use in our study. While random mutations in SMILES (especially those involving special symbols) often result in invalid molecules, the SELFIES grammar ensures that all generated sequences decode to valid molecules. (Figure inspired by [32].)

**Table 1:**
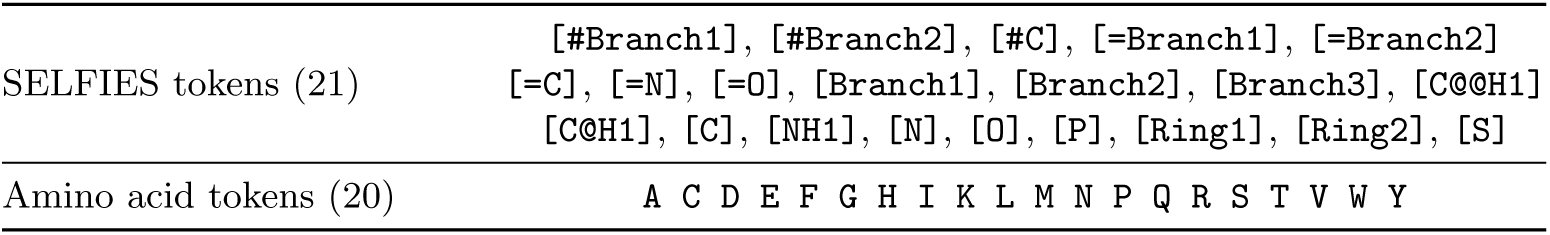
SELFIES tokens used in this research compared to the 20 canonical amino acid tokens.

|  |  |
| --- | --- |
| SELFIES tokens (21) | [#Branch1], [#Branch2], [#C], [=Branch1], [=Branch2] |
|  | [=C], [=N], [=O], [Branch1], [Branch2], [Branch3], [C@@H1] |
|  | [C@H1], [C], [NH1], [N], [O], [P], [Ring1], [Ring2], [S] |
| Amino acid tokens (20) | A C D E F G H I K L M N P Q R S T V W Y |

### 2.3 Discrete Diffusion Model and Denoising Architecture

We opted to use the discrete denoising diffusion probabilistic model (D3PM) framework due to its competitive performance and the flexibility offered by interchangeable transition probabilities. Furthermore, D3PM enables parallel generation by denoising all sequence positions simultaneously at each step. To model the conditional probability of the reverse process, *p_θ_*(*x*^_0_|*x_t_*), we chose the ByteNet neural network architecture [42], which demonstrated promising results in protein sequence design for EvoDiff [19], particularly in handling long sequences efficiently and capturing long-range dependencies. More details on the ByteNet architecture can be found in Supplementary Material A.2.

#### 2.3.1 Discrete Denoising Diffusion Probabilistic Models (D3PMs)

D3PM [18] is a discretized generalized version of DDPM [14] (details in Supplementary Material A.3). In D3PM, the forward process for a scalar random variable with *K* categories (*x_t_, x_t_*_−1_ ∈ 1*, …, K*) is defined by a probabilistic transition matrix, represented as [***Q****_t_*]*_ij_* = *q*(*x_t_* = *j* | *x_t_*_−1_ = *i*). When we denote the row vector ***x*** as its one-hot version, we can write

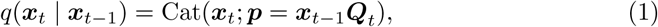

where Cat(***x***; ***p***) is a categorical distribution over the one-hot row vector ***x*** with probabilities given by the row vector ***p***, and ***x****_t_*_−1_***Q****_t_* is a row vector-matrix product. From this notation, we derive the two criteria necessary for a noise distribution in a diffusion process:

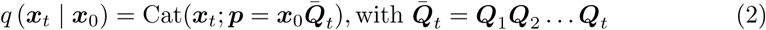

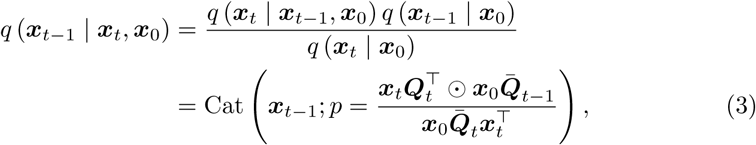

here, Equation 2 shows how the noise for any timestep *t* can be efficiently calculated. Equation 3 describes how the tractable forward posterior can be computed using Bayes’ rule. Using this approach, we can set the transition matrices to any noise schedule. In our study, we focus on two noise schedules (see Table 2): uniform noise, which randomly replaces each token with any other token with equal probability, and absorbing (or masking) noise, where tokens progressively transition to a fixed absorbing state (typically denoted as #). Examples of a uniform and absorbing transition matrix for a random variable with three categories at an arbitrary timestep *t* are shown below. The uniform matrix (Equation 4) consists of the three core categories with equal transition probabilities to other categories. The absorbing matrix (Equation 5) includes a fourth, masked category that acts as an absorbing state—once entered, all transitions lead exclusively to this category. For both uniform and absorbing noise schedules, we can set the noising parameter to *β_t_* = (*T* − *t* + 1)^−1^ as given by the original D3PM [18]. As *t* increases, *β_t_* increases, and the matrix converges to the specified noise distribution.

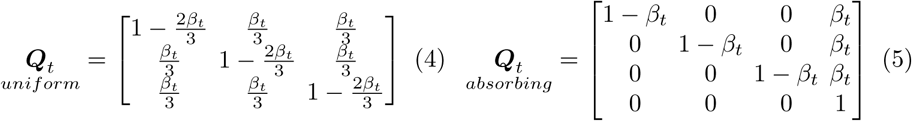

**Table 2:**
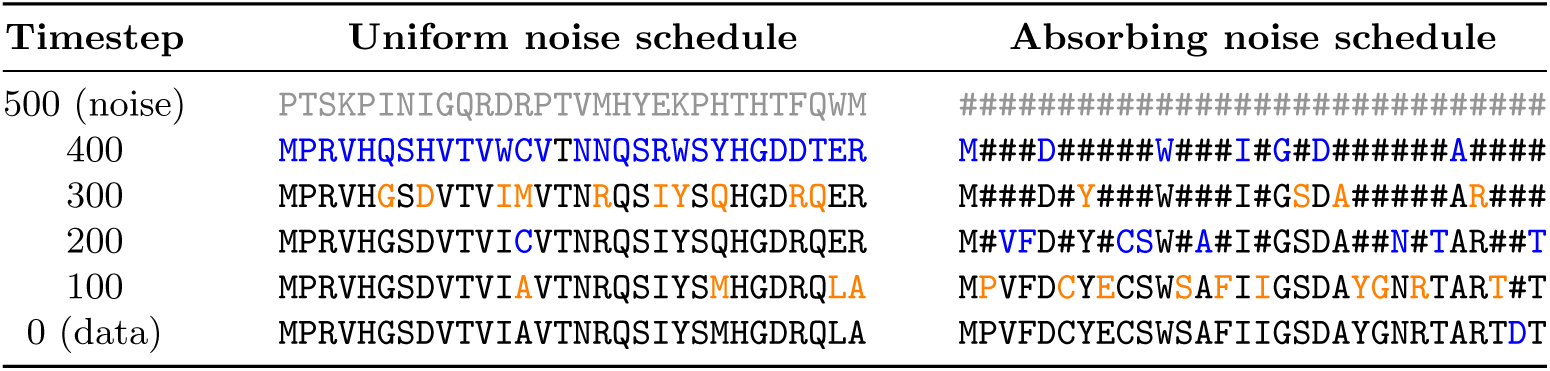
Sequence generation progression for the amino acid representation using both uniform and absorbing noise schedules. Tokens updated at each timestep relative to the previous one are highlighted in alternating blue and orange colors. The absorbing noise process only allows changes from the absorbing state to an amino acid token.

Lastly, an updated loss function is integrated into the diffusion model. Austin et al. [18] use an alternative hybrid loss function, which leads to improved quality of samples:

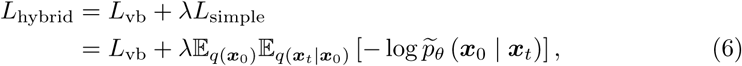

where they introduce an extra denoising objective for the ***x***_0_-parametrization of the reverse process, that encourages good predictions of the data ***x***_0_ at each timestep. This additional objective corresponds to the cross-entropy term *L*_0_ (at *t* = 1) in DDPM (Supplementary Material A.3), weighted by the *λ* parameter.

#### 2.3.2 Training and Generation

During the D3PM training phase, the denoising model takes as input a protein sequence and a diffusion timestep. Protein sequences are tokenized according to the selected representation (either SELFIES or amino acid) and mapped to embedding vectors. The diffusion timesteps *t* are encoded into vectors of the same dimension using sinusoidal positional encoding. The sequence embeddings and diffusion timestep encodings are added element-wise and then fed through several ByteNet blocks. The output of the last ByteNet block is embedded back into the protein sequence representation space using a linear layer.

During the inference or sequence generation process, we start with a fully noised sequence given by the noise schedule. We give this sequence and the timestep *T* as input to our model. We then iteratively, predict *p_θ_*(***x̂***_0_|***x****_t_*) and calculate its posterior *q* (***x****_t_*_−1_ | ***x****_t_,* ***x̂***_0_). Using this posterior, we sample the next timestep ***x****_t_*_−1_ from a multi-nomial distribution and feed it into our model together with timestep *t* − 1. After progressing through all timesteps, we end up with ***x̂***_0_. Examples of the generation process on both amino acid and SELFIES sequences for the two noise schedules can be found in Table 2 and Supplementary Material C.

### 2.4 Proposed All-Atom-Level Evaluation and Filtering

The models trained using the all-atom SELFIES representation focus exclusively on canonical proteins (from UniRef50). However, our experiments revealed that the outputs are not always valid proteins and often include residues beyond the 20 canonical amino acids. While SELFIES ensures chemical validity, this alone does not guarantee that the generated molecules are structurally consistent with real proteins. To assess how well the all-atom model generates protein-like molecules, we developed a set of metrics that evaluate the presence of peptide bonds and a continuous backbone, as well as the constitutional and stereochemistry correctness of the generated amino acids, which we use to construct the amino acid sequence from the SELFIES output. This is the first part of the full evaluation workflow for the all-atom-generated sequences, which is shown in Fig. 2.

**Fig. 2:**
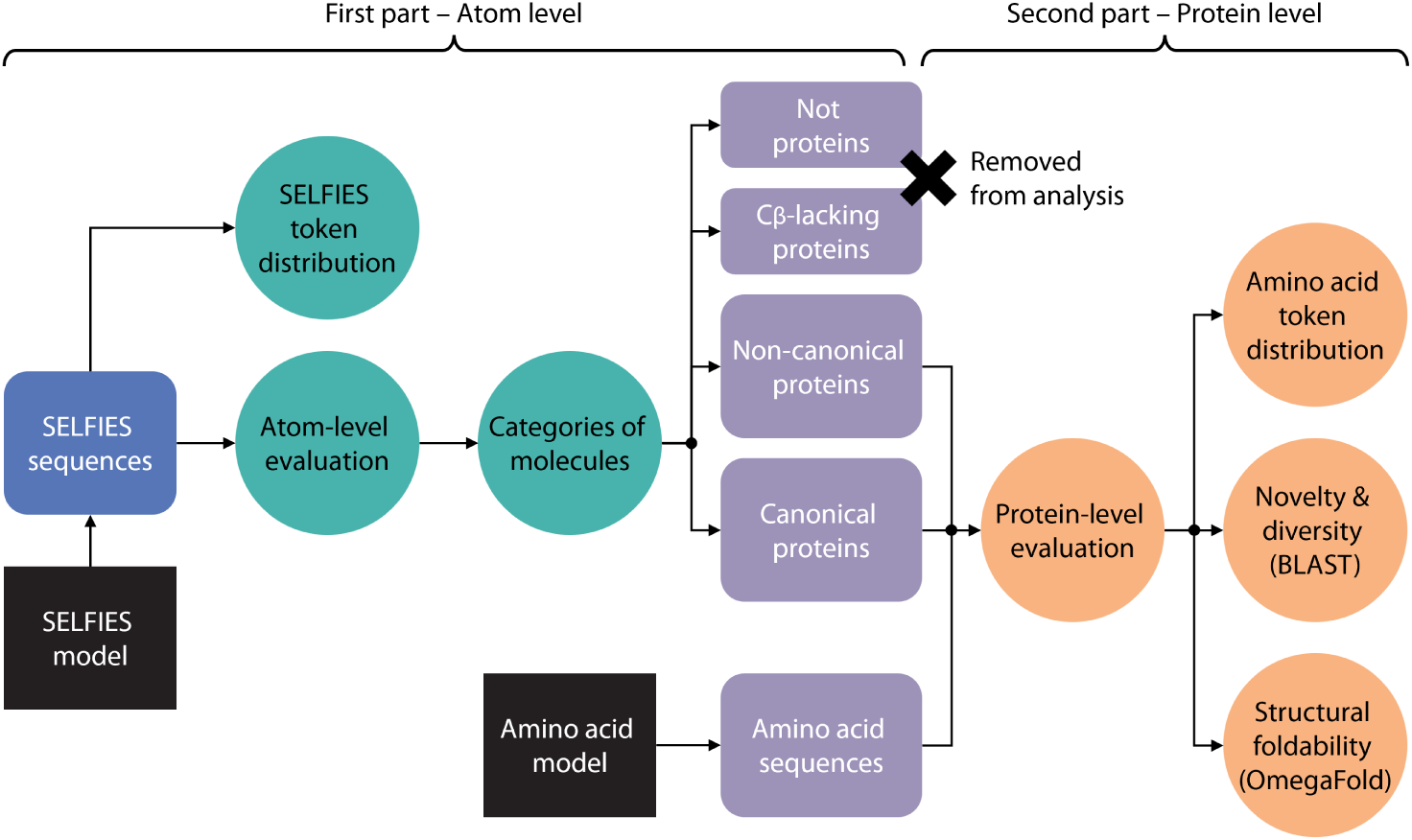
Evaluation workflow for the SELFIES and amino acid generated sequences. All-atom SELFIES sequences are analyzed through various stages, including atom-level evaluation and molecule categorization (in green color), as well as protein-level evaluation of filtered non-canonical and canonical protein sequences (in orange color).

#### Categorization of generated molecules

We categorize the generated molecules into the following four classes:

- **Not a protein**: A molecule in which a continuous backbone cannot be constructed or no peptide bonds are present. Such molecules fail to meet the fundamental structural criteria of a protein.
- *C_β_***-lacking protein**: A protein where at least one amino acid side chain (excluding Glycine) does not begin with a carbon (i.e. lacks a *β*-carbon atom). This deviates from both standard amino acid structures and most non-canonical amino acids.
- **Non-canonical protein**: A protein containing non-canonical amino acids or amino acids that are only constitutionally correct but not stereochemically correct. Additionally, molecules failing the SMILES check—indicating extra atoms at the beginning or end of the backbone—are classified as non-canonical.
- **Canonical protein**: A protein composed exclusively of canonical amino acids with correct stereochemistry that passes the SMILES check. This indicates that the generated molecule exhibits all the traits of a canonical protein constructed solely from natural amino acids.

We outline each step of our evaluation pipeline and the corresponding validation criteria below.

#### Step 1: Identify peptide bonds and a continuous backbone

The first step in our analysis involves converting each generated SELFIES sequence into a SMILES string. Since SELFIES guarantees chemical validity, the resulting SMILES also corresponds to a valid molecule. However, the original SELFIES may contain extra tokens that are ignored during decoding. To quantify these unused tokens, we perform a round-trip conversion: each SELFIES sequence is converted to SMILES and then back to SELFIES. Because SMILES only retains atoms and bonds that contribute to a valid structure, this reciprocal check reveals how many tokens in the original SELFIES sequence were effectively discarded.

The SMILES sequence is then transformed into an RDKit [40] molecule graph object for detailed examination. Our evaluation method relies on performing substructure searches within the molecule and conducting graph traversals of the side chains. Since certain amino acids are substructures of others, directly searching for complete amino acids is ineffective. We begin by *identifying peptide bonds*^1^ through substructure searches. Identifying peptide bonds allows us to locate a *continuous backbone* within the molecule. In Fig. 3 we can see a generic amino acid, a peptide bond in a continuous backbone, and two examples of graph traversal on the side chain.

**Fig. 3:**
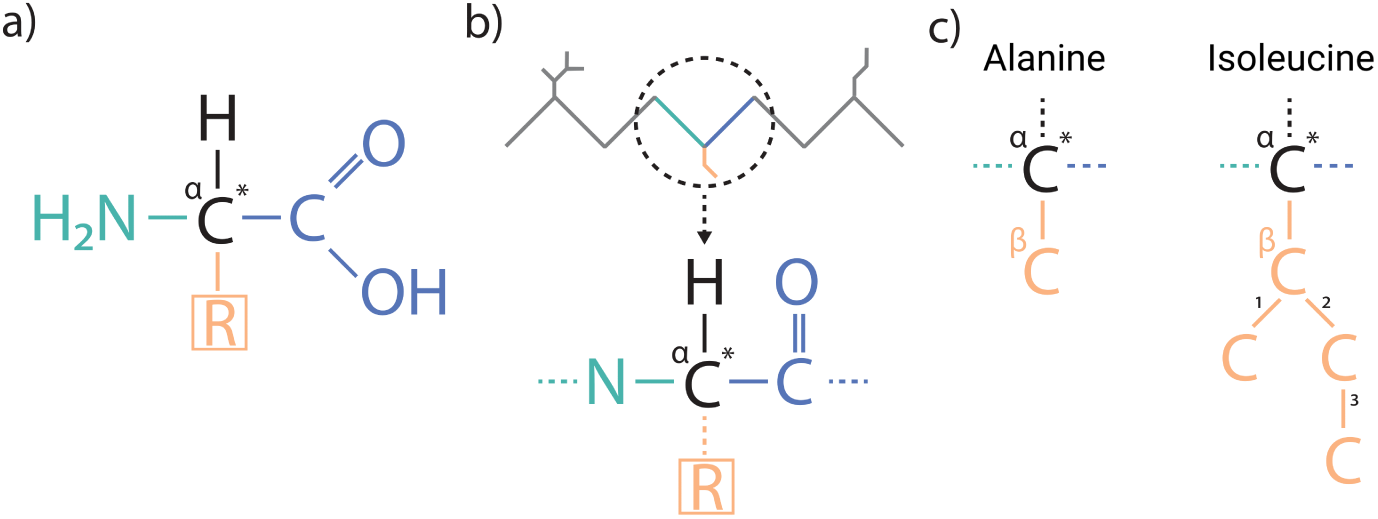
(a) The molecular structure of an amino acid in its non-ionized form, showing the central *α*-carbon (black), the carboxyl group (blue), the amino group (green), and the variable side chain (orange). The asterisk marks the chiral center. **(b)** A section of a protein backbone highlighting a peptide bond, with the *α*-carbon indicated as a reference point for side chain analysis. **(c)** Two examples of side-chain structures with *α*-*β* carbon bonds: Alanine and Isoleucine. The numbers illustrate the graph traversal order over the side chain bonds, which is used to analyze the molecular structure of the side chain during evaluation. Notably, Alanine can be seen as a substructure of Isoleucine.

#### Step 2: Analyze side-chains, presence of beta-carbon atom, constitutionally and stereochemistry correctness

Given a continuous backbone and its peptide bonds, we analyze the side chains attached to each *α*-carbon atom. From the *α*-carbon, we find the beginning of the side chain. If no atoms are found, we classify that peptide bond as Glycine, the only amino acid without a side chain. If a side chain is present, we observe the first atom; if it is a carbon atom, we have identified an *α*-*β* carbon bond. Using the RDKit molecule object, we then perform a breadth-first search (BFS) graph traversal of the side chain, exploring new bonds not already part of the backbone or the side chain. If we find a side chain that does not start with a carbon atom, we mark that residue as *β-carbon lacking* and do not analyze the side chain further. Otherwise, if the atomic structure of the side chain matches one of the 20 natural amino acids, we mark the residue as *constitutionally correct*.

Next, we check for the *stereochemistry correctness* of the generated amino acids. All amino acids, except Glycine, have a chiral center, meaning they can exist in two forms [43]. These forms, called stereoisomers, have the same molecular composition but their spatial 3D structures are mirror images of each other, flipped around the chiral center. Although the two stereoisomers exhibit identical physical and chemical properties, we check for amino acids in the *L*-form, as these are the ones found in living organisms.

#### Step 3: Build canonical and non-canonical proteins

If an amino acid meets both constitutional and stereochemistry correctness criteria, we classify it as canonical; otherwise, it is considered non-canonical. We then construct an amino acid sequence for the continuous backbone. Since each side chain was matched to one of the 20 natural amino acids in Step 2, we record canonical residues using their standard one-letter codes and mark non-canonical ones with X (unknown). Finally, we translate the amino acid sequence back into a SMILES representation. By comparing this string to the original SMILES (from the generated SELFIES), we can evaluate whether the model introduced extra atoms at the beginning or end of the backbone (*SMILES check*).

The outcome of this process is the classification of generated molecules into the four categories defined above (and illustrated in Fig. 2). In summary, Step 1 identifies molecules that are not proteins, Step 2 identifies proteins lacking *C_β_* atoms in their side chains, and Steps 2 and 3 together help classify valid proteins into non-canonical or canonical. By applying this evaluation pipeline, we assess how effectively the all-atom model generates valid proteins and identify where errors occur.

### 2.5 Protein-Level Evaluation

We then assess the quality of the generated amino acid sequences using sequence similarity metrics to evaluate novelty and diversity, and 3D structure prediction confidence to estimate structural foldability (second part in Fig. 2).

#### Novelty and diversity

To assess the uniqueness and variability of the generated sequences, we use two metrics: *novelty*, which measures the generation of sequences distinct from the training data, and *diversity*, which quantifies the variation within the generated set.

We use BLAST (basic local alignment search tool) [44] to identify regions of similarity between protein sequences. In BLAST, unknown amino acids (X) are treated as ambiguous residues that can align with any amino acid but do not contribute to the overall alignment score. For each protein search, we record the number of matches, the number of unique matches, and key alignment metrics including the e-value, score, query coverage, and percentage identity. A match is considered significant if its e-value falls below a predefined threshold. For each generated protein set, we also compute the proportion of sequences with at least one significant match. Based on this, we define *novelty* as the proportion of generated sequences with no significant matches to the training dataset, and *diversity* as the proportion of sequences with no significant matches to any other sequence within the generated set. Our initial experiments using a strict e-value threshold of 10^−5^ found no significant matches (Supplementary Fig. S4). Therefore, we opted for a threshold of 0.05 to balance between sensitivity and specificity in detecting significant alignments.

#### Structure foldability

We use the structure predictor OmegaFold [45] to evaluate the *structure foldability* of the generated protein sequences. OmegaFold provides a confidence score called pLDDT (predicted local distance difference test) for each residue in the protein, ranging from 0 to 100. In our evaluation, we use pLDDT confidence scores averaged across the whole sequence to assess the structural soundness of the generated proteins. Since OmegaFold does not predict atomic coordinates for unknown amino acids (X), these residues are excluded from the pLDDT calculation. However, their sequence positions are considered, making them appear as gaps in the predicted structure. We consider sequences with average pLDDT scores above 70 reliable and below 50 to be unreliably predicted, following established conventions in protein structure prediction [7]. Scores between 50 and 70 indicate moderate confidence, suggesting the predicted structures may have some correct regions but also areas of high uncertainty.

### 2.6 Experimental Setup

#### Implementation and training

We used the 38M-parameter ByteNet model from EvoDiff. We adapted the original PyTorch implementation to incorporate the SELFIES tokenization and the absorbing noise schedule in the diffusion process. We trained four discrete diffusion models: SELFIES and amino acid representations, uniform and absorbing noise processes. Each model was trained on the UniRef50 dataset with the same architecture and hyperparameters (listed in Supplementary Table S7), using a single NVIDIA A40 GPU. Due to the longer sequence lengths in the SELFIES representation, the total number of tokens in the training set is significantly higher than for the amino acid representation. As a result, the SELFIES models are exposed to more tokens per epoch and converge in fewer epochs (8 epochs, 312 hours), while the amino acid models required 30 epochs (208 hours) to achieve comparable convergence. Training and validation curves can be found in Supplementary Figures S9 and S10.

#### Evaluation

We first evaluated the performance of the all-atom SELFIES models on atom-level metrics for protein filtering and categorization. We generated 1000 sequences of random lengths between the minimum and maximum lengths from the training set (225 to 1907 SELFIES tokens). We refer to this set as *SELFIES (unfiltered)*. We also generated 1000 sequences from the amino acid models (lengths between 30 and 100 amino acids). To ensure a fair comparison, we generated SELFIES sequences until we obtained 1000 non-canonical and 1000 canonical valid proteins. These proteins were filtered to lengths between 10 and 120 amino acids, with no more than 4 non-canonical residues (denoted as unknown X). We refer to these three sets with valid proteins as i. *Amino acid*, ii. *SELFIES (non-canonical)*, and iii. *SELFIES (canonical)*. We then conducted protein-level evaluations using novelty, diversity, and structure confidence as metrics. We additionally run experiments to evaluate the performance across different sequence lengths. This analysis reveals how increasing sequence length affects the models’ ability to generate valid and structurally sound proteins.

## 3 Results and Discussion

Our study shows the feasibility of generating valid protein sequences using an all-atom SELFIES representation within discrete diffusion models, marking a step forward in protein design methodologies. In this section, we analyze the performance of the models using both atom-level and protein-level metrics and assess the novelty, diversity, and structural foldability of the generated sequences.

### 3.1 Absorbing Noise Enhances Validity of SELFIES-Generated Proteins

We first assess the performance of the all-atom SELFIES models using our proposed atom-level evaluation pipeline. Our analysis reveals that the absorbing model consistently outperforms the uniform model in generating more valid protein sequences.

As shown in Fig. 4, both models generate distributions of SELFIES tokens that closely resemble the training set, indicating successful learning of the all-atom vocabulary. However, when we analyze how often pairs of SELFIES tokens appear together (2-mer distribution), only 291 out of 441 possible pairs are found in the training set. In contrast, the uniform model generates all 441 pairs, with a KL divergence from the training distribution of 0.034. The absorbing model generates slightly fewer combinations (426), which more closely match the training distribution, with a KL divergence of 0.007. This improved alignment also translates into better performance: the absorbing model outperforms the uniform model by producing sequences with fewer unused tokens, fully continuous backbones, and a higher proportion of constitutionally and stereochemically correct amino acids (Table 3).

**Fig. 4:**
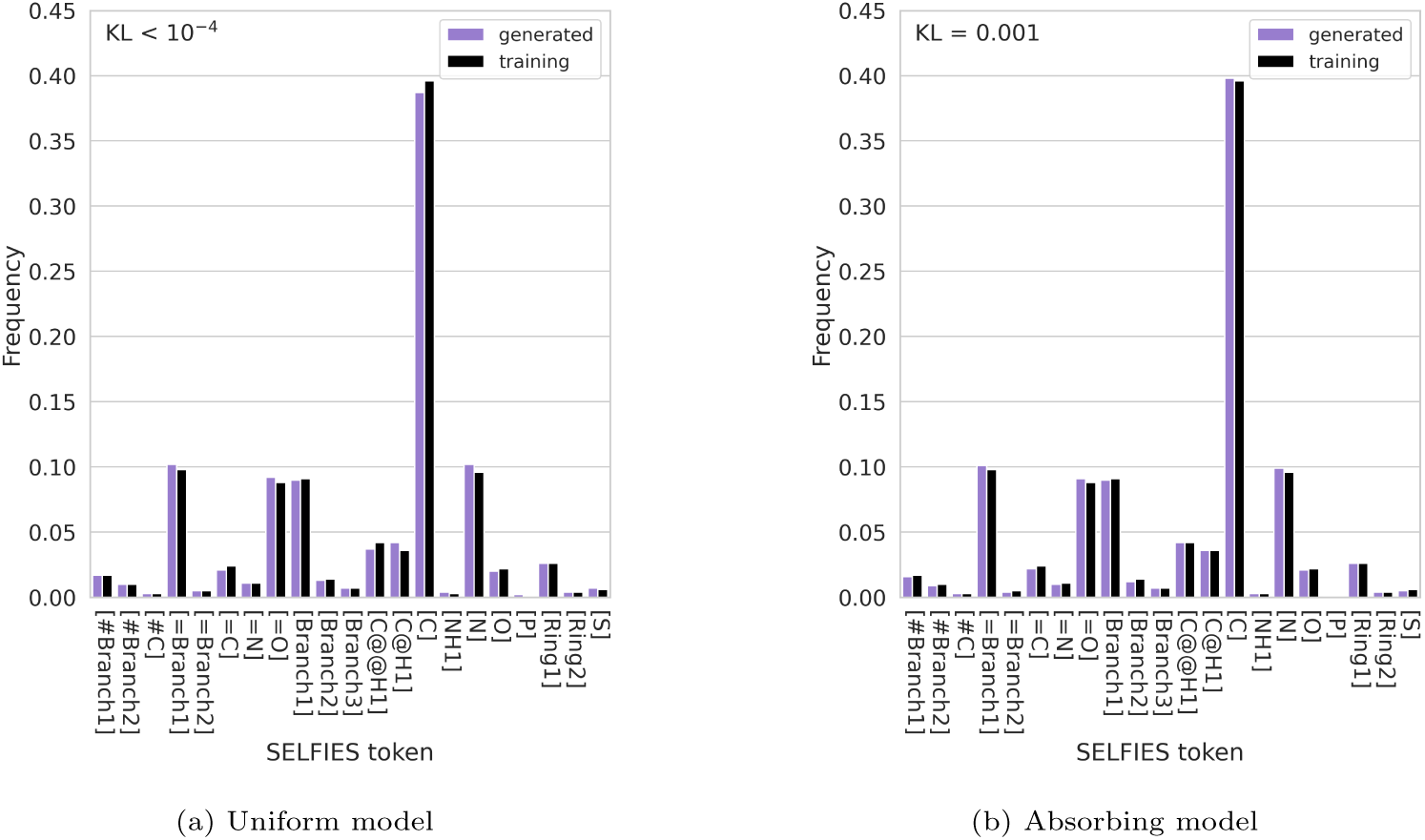
SELFIES token distributions for the 1000 unfiltered all-atom sequences generated by the **(a)** uniform and **(b)** absorbing noise models. Token frequency is calculated as the count of each token divided by the total number of tokens across all generated proteins. We compare to the training set distribution (black bars) and report the Kullback–Leibler divergence, KL(*P_training_*||*Q_generated_*). Both models closely match the training distribution. Notably, the carbon token [C] is the most abundant across all distributions, consistent with organic chemistry.

**Table 3:**
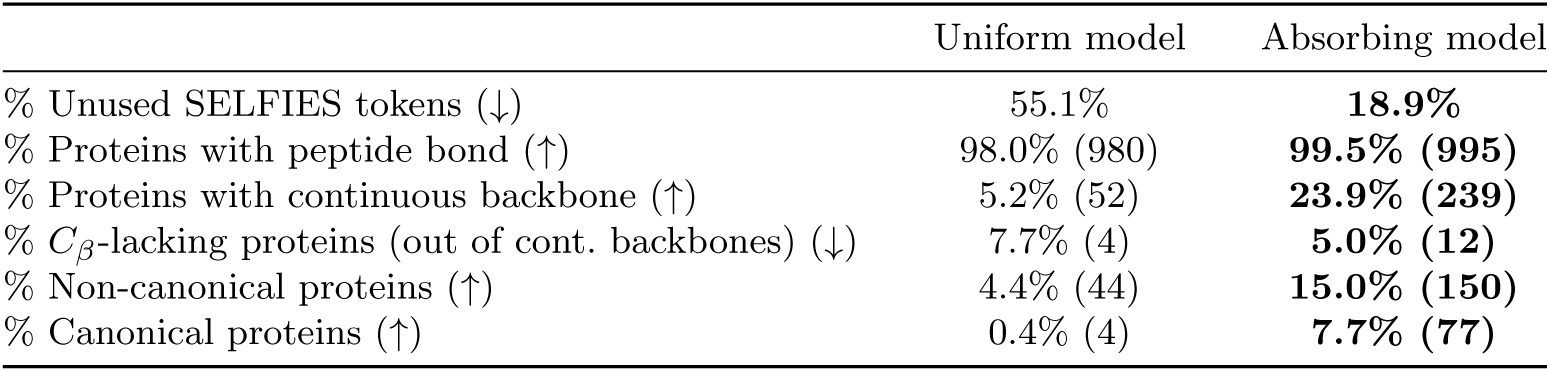
Unused SELFIES tokens and protein categorization for the 1000 unfiltered all-atom SELFIES sequences generated by the uniform and absorbing noise models. Best results are highlighted in bold.

Table 3 shows that the absorbing model generates 239, 093 unused SELFIES tokens (18.9% of the total), compared to 686, 273 (55.1%) from the uniform model. The higher unused token ratio observed in the uniform model likely results from its noise process, which allows for token alterations late in the generation process, potentially disrupting sequence coherence. In contrast, the absorbing model does not allow changes to already unmasked tokens, preserving earlier sequence decisions. Moreover, while both models generate sequences with at least one peptide bond—a fundamental feature of proteins—the absorbing model outperforms the uniform model in producing fully continuous backbones (239 compared to 52, out of 1000). This difference highlights the absorbing model’s ability to make fewer errors by using the SELFIES token space more efficiently and maintaining chemical integrity throughout the sequence generation process.

In addition, the absorbing model generates more non-canonical and canonical proteins validated by SMILES conversion: 150 and 77, compared to only 44 and 4 from the uniform model. The absorbing model also produces 12*/*239 (5.0%) *C_β_*-lacking proteins with continuous backbones, compared to 4*/*52 (7.7%) in the uniform model. These relatively low occurrences suggest that both models primarily generate sequences with correct amino acid side chains.

Out of all the generated proteins with continuous backbones (Table 4), the absorbing model produces 12, 647 constitutionally correct canonical amino acids, compared to 1180 for the uniform model—a tenfold increase. This suggests that the absorbing model better captures amino acid-level features. While over 90% of amino acids in these sequences are constitutionally correct, the low total count for the uniform model indicates that the generated proteins are very short, as reflected in the relatively low sequence lengths of the non-canonical and canonical proteins (30 amino acids or less on average). Moreover, for both models, the generated canonical amino acids are mostly stereochemically correct, showcasing the models’ ability to capture the stereochemical properties of the dataset.

**Table 4:** Amino acid-level metrics for all-atom SELFIES sequences with continuous backbones, generated by the uniform and absorbing noise models. Average per-sequence ratios are reported in parentheses. The average sequence lengths (in amino acids) are also provided for both non-canonical and canonical protein sets. Best results are highlighted in bold.

|  | Uniform model | Absorbing model |
| --- | --- | --- |
| N. Non-canonical amino acids (↑) | 93 (8.9%) | **276 (1.9%)** |
| N. Constitutionally correct amino acids (↑) | 1180 (90.7%) | **12647 (98.0%)** |
| N. Stereochemistry correct amino acids (↑) | 1168 (89.9%) | **12634 (97.9%)** |
| Non-canonical protein sequence length (avg.±std. ↑) | 24.1 ± 13.3 | **60.9 ± 30.7** |
| Canonical protein sequence length (avg.±std. ↑) | 30.5 ± 7.2 | **37.3 ± 15.3** |

Next, we examine the results across different SELFIES sequence lengths (number of tokens), as shown in Supplementary Table S3. While both models consistently produce at least one peptide bond per sequence across all length ranges, the number of detected continuous backbones, non-canonical proteins, and canonical proteins decreases as the SELFIES length increases. For the uniform model, the percentage of unused SELFIES tokens increases sharply. This is consistent with the model’s difficulty in producing proteins longer than 40 amino acids on average (Table 4 and Supplementary Table S3), despite being trained on natural proteins up to 100 amino acids. In contrast, the absorbing model shows a more gradual increase in unused SELFIES tokens with sequence length, and it can generate non-canonical proteins even for the longest sequence lengths tested. Notably, the absorbing model produces non-canonical proteins with lengths ranging from 26 to 138 amino acids on average (see Supplementary Table S3), extending beyond the maximum protein sequence length of 100 on which it was trained. This indicates that the absorbing noise process helps generalize to longer valid sequences.

### 3.2 Absorbing Models More Accurately Reflect Amino Acid Token Distributions

We now compare the 1000 sequences generated by the amino acid models (using uniform and absorbing noise schedules) with non-canonical and canonical protein sequences filtered from the SELFIES models. To ensure a fair comparison, we continue generating SELFIES sequences until obtaining 1000 valid proteins of each type. Achieving this required generating 160, 000 SELFIES sequences with varying lengths with the uniform model and 20, 000 with the absorbing model, re-emphasizing the uniform model’s limited ability to produce fully valid proteins. The resulting amino acid sequence length distributions for each set and noise schedule are shown in Supplementary Fig. S6. The SELFIES models generate proteins outside the range of 30 to 100 amino acids, due to the all-atom representation and the lack of direct control over protein sequence length from the SELFIES strings. Moreover, as previously observed, the SELFIES uniform model tends to generate shorter proteins than the absorbing model, and is unable to produce sequences longer than 70 amino acids.

The amino acid token distributions for the three sets are illustrated in Fig. 5. The amino acid models, particularly the absorbing one, produce token distributions that closely match the training set, effectively capturing the natural amino acid composition. In contrast, valid proteins generated by the SELFIES models show some deviation from the training distribution; however, these differences remain relatively minor, as reflected by their low Kullback-Leibler (KL) divergence values. In particular, greater deviations—and correspondingly higher KL values—are expected for the non-canonical proteins due to the inclusion of an additional X token, which represents unknown amino acids.

**Fig. 5:**
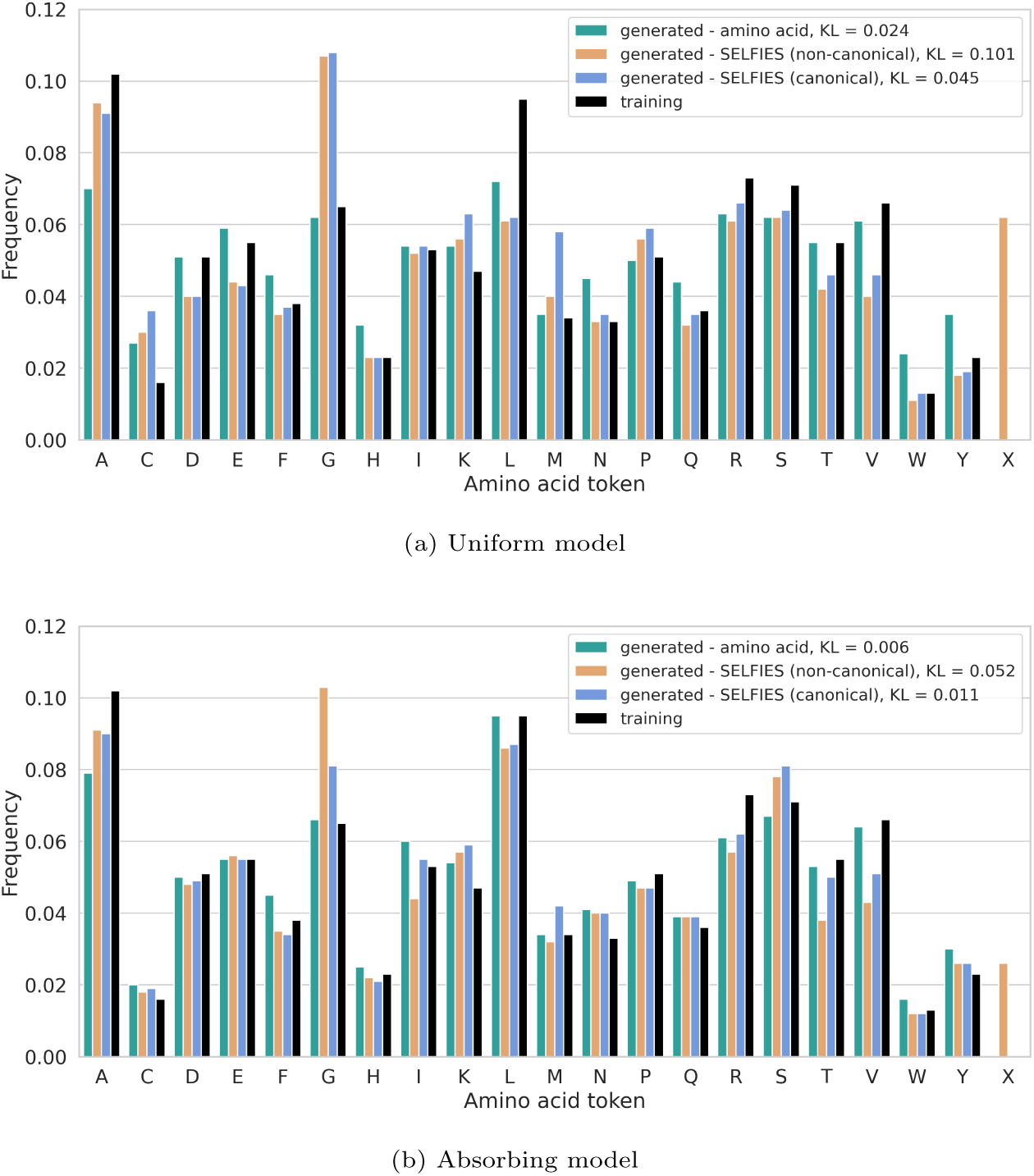
Amino acid token distributions for the 1000 valid proteins in each of the amino acid, SELFIES non-canonical, and SELFIES canonical sets, generated by the **(a)** uniform and **(b)** absorbing noise models. Within each generated set, token frequency is calculated as the count of each token divided by the total number of tokens across all proteins. The X token denotes unknown non-canonical amino acids found exclusively in the generated SELFIES non-canonical proteins. We compare to the training set distribution (black bars) and report the Kullback–Leibler divergence, KL(*Ptraining* ||*Qgenerated*).

Interestingly, most SELFIES models tend to generate a considerably higher proportion of Glycine (G), the simplest amino acid with no side chain—though this effect is less pronounced in the SELFIES absorbing canonical set. This suggests a bias toward chemically simpler structures, either during generation or as a result of post-processing filters. Conversely, the models underrepresent the amino acids Threonine (T) and Valine (V), despite their side chains not being particularly long. Threonine, in particular, has a branched side chain with a hydroxyl group, introducing both polarity and stereochemical complexity that may make it more challenging to generate correctly.

Across all uniform models, including the amino acid one, we also observe a consistently higher frequency of Cysteine (C) and a lower frequency of Leucine (L), despite Leucine being well represented in the training set. Leucine is chemically similar to Valine, with both having branched nonpolar side chains (Leucine being slightly longer by one methylene group). The lower frequency of both Valine and Leucine may reflect challenges in modeling branched, hydrophobic side chains. In contrast, the elevated occurrence of Cysteine may be due to its simple side chain and distinctive sulfur atom, making it easier for the models to identify or favor during generation.

### 3.3 SELFIES Models Generate Highly Novel and Diverse Sequences

Next, we assess the novelty and diversity of sequences generated by the SELFIES models compared to the amino acid models. Overall, the BLAST results indicate that the SELFIES models produce a higher proportion of novel and diverse sequences. Table 5 summarizes these results, reporting the percentage of sequences with no significant matches (e-value below 0.05). Notably, across all models and representations, more than 94% generated sequences lack significant matches, demonstrating consistently high novelty and diversity.

**Table 5:** Percentage of **(a)** novel (e-value *<* 0.05 to the training set) and **(b)** diverse (e-value *<* 0.05 within the generated set) sequences based on BLAST analysis. Results are shown for the uniform and absorbing models, comparing the 1000 valid proteins in each of the amino acid, SELFIES non-canonical, and SELFIES canonical sets. Higher is better for both metrics. Best results are highlighted in bold.

| (a) Novelty |  |  | (b) Diversity |  |  |
| --- | --- | --- | --- | --- | --- |
| Protein set | Uniform model | Absorbing model | Protein set | Uniform model | Absorbing model |
| Amino acid | 95.7% | 94.6% | Amino acid | 94.6% | 95.1% |
| SELFIES (non-canonical) | **99.9%** | 97.8% | SELFIES (non-canonical) | **99.7%** | 98.7% |
| SELFIES (canonical) | 99.8% | 98.5% | SELFIES (canonical) | 97.8% | 96.4% |

The novelty results (Table 5a) show that proteins generated by the SELFIES models—both non-canonical and canonical—consistently exhibit higher novelty than those produced by the amino acid models. Specifically, the SELFIES uniform model achieves the highest scores, with 99.9% of non-canonical and 99.8% of canonical sequences showing no significant matches to the training set, indicating that nearly all generated proteins are novel. The absorbing SELFIES model also performs well, though with slightly lower novelty. In contrast, the amino acid models yield lower novelty, with a maximum of 95.7% novel sequences for the uniform model, suggesting higher similarity to training data. A similar trend is observed for diversity (Table 5b). The SELFIES uniform model also outperforms, producing highly diverse protein sets: 99.7% of non-canonical and 97.8% of canonical proteins show no significant similarity to other sequences within each generated set. This indicates high sequence variability across the generated proteins. The SELFIES absorbing model also performs well, though diversity among canonical proteins (96.4%) is more comparable to that of the amino acid absorbing model (95.1%).

Additionally, when evaluating sequence similarity based on BLAST score, query coverage, and percentage identity (Supplementary Table S5), these metric values are consistently lower for diversity than for novelty. This indicates that, while more matches occur within the generated sets, the sequences are less similar to each other than to those in the training set, which also favors diversity.

### 3.4 SELFIES Absorbing Model Shows Enhanced Structural Foldability Across a Wider Range of Sequence Lengths

We further investigate the structural foldability of the generated proteins by predicting their 3D structures with OmegaFold. Our results indicate that, while the all-atom SELFIES representation enhances novelty and diversity, this may come at the cost of structural confidence, particularly for longer sequences. Achieving a balance between these properties is crucial for protein design, as highly novel sequences risk being non-foldable structures, compromising their functionality.

Table 6a shows that, on average, the all-atom SELFIES representation contributes to generating proteins with better predicted structures. Specifically, the canonical proteins generated by the SELFIES uniform model achieve the highest mean pLDDT score of 64.2. Although all models have mean pLDDT scores above 50, none reach the threshold of 70, which is considered indicative of reliable protein folding. However, when examining the percentage of proteins with pLDDT scores above 70 (Table 6b), the SELFIES uniform model generates more foldable proteins than the amino acid models: 32.1% of proteins from the SELFIES canonical set are foldable. Additionally, Supplementary Fig. S7 illustrates the pLDDT score distributions for each model and representation. Notably, almost all generated distributions from the SELFIES models have lower standard deviations than the amino acid models and exhibit higher proportions of foldable proteins (pLDDT *>* 70). While these results might suggest a limitation of the models in generating reliably foldable proteins, it is important to consider that the training data also plays a significant role. We observe that even a subset of 100k training proteins does not reach the pLDDT threshold of 70 on average (see distributions in Supplementary Fig. S8). Since many sequences in the training set lack experimentally resolved structures, structure predictors like OmegaFold may not accurately assess their foldability—an issue that could also affect the evaluation of generated sequences.

**Table 6:** Structural foldability measured as OmegaFold average pLDDT confidence scores. Results are shown for the uniform and absorbing models, comparing the 1000 valid proteins in each of the amino acid, SELFIES non-canonical, and SELFIES canonical sets. For each set, we report **(a)** the average of all pLDDT scores and **(b)** the percentage of proteins with pLDDT greater than 70 (indicating reliable folding). We also compare with the pLDDT score of 100k samples from the training set. Higher is better for the pLDDT score. Best results are highlighted in bold.

| (a) Average pLDDT score |  |  | (b) % Proteins with pLDDT > 70 |  |  |
| --- | --- | --- | --- | --- | --- |
| Protein set | Uniform model | Absorbing model | Protein set | Uniform model | Absorbing model |
| Amino acid | 53.7 ± 12.7 | 58.5 ± 13.1 | Amino acid | 11.0% | 19.4% |
| SELFIES (non-canonical) | 62.3 ± 12.3 | 57.0 ± 12.7 | SELFIES (non-canonical) | 27.2% | 16.2% |
| SELFIES (canonical) | **64.2 ± 11.9** | 62.7 ± 12.5 | SELFIES (canonical) | **32.1%** | 27.9% |
| Train (100k) | 66.9 ± 15.3 | | Train (100k) | 43.6% | |

When examining pLDDT results for different amino acid sequence lengths (Fig. 6, and Supplementary Table S6 for numerical details), we observe a clear trend: longer protein sequences exhibit lower average pLDDT scores. This is expected, as longer sequences are more complex and prone to cumulative errors during generation, leading to less accurate structures. Furthermore, when we exclude protein sequences shorter than 30 amino acids, the amino acid absorbing model consistently achieves higher average pLDDT scores across the different protein length groups. This indicates that the overall pLDDT results for the SELFIES models are mostly influenced by shorter sequences (less than 30 amino acids) with higher mean pLDDT scores. In contrast, the amino acid models were not tasked with generating sequences below 30, resulting in lower means overall. Despite this, the SELFIES absorbing model generates protein sequences across a wide range of amino acid lengths. It achieves the highest mean pLDDT score (68.3) for short canonical proteins (10 to 30 amino acids), approaching the 70 threshold. However, for longer sequences, the amino acid models perform better and yield a higher proportion of foldable proteins (pLDDT *>* 70), especially the absorbing model. While the SELFIES absorbing model exhibits slightly weaker performance in terms of structural confidence, it remains a preferred option for applications requiring a broader range of sequence lengths.

**Fig. 6:**
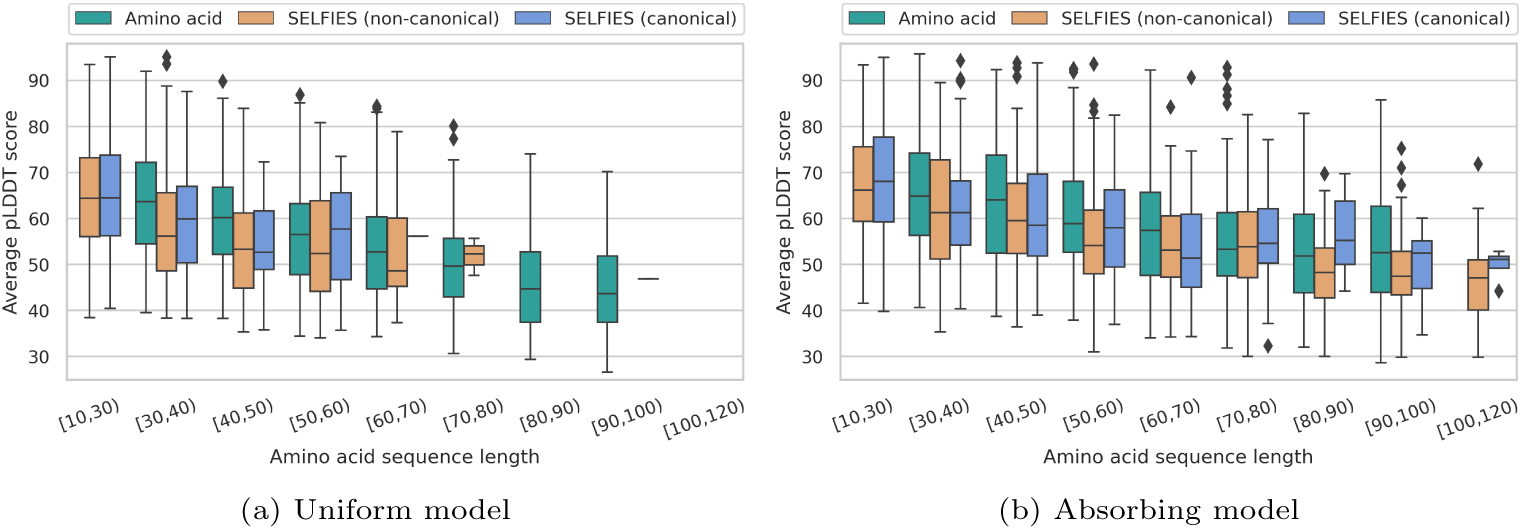
Per-sequence length distribution of average pLDDT confidence scores for the 1000 valid proteins in each of the amino acid, SELFIES non-canonical, and SELFIES canonical sets, generated by the **(a)** uniform and **(b)** absorbing noise models. pLDDT scores are shown in length ranges from 10 to 120 amino acids.

## 4 Conclusion

Our work illustrates the significant potential of discrete diffusion models using, for the first time, an all-atom representation for protein sequence generation. This approach offers a more detailed and flexible framework compared to traditional amino acid representations, enabling the incorporation of non-canonical amino acids and posttranslational modifications.

While our study primarily focuses on natural protein sequences, we also observe the generation of non-canonical amino acids in the SELFIES models. This opens an avenue for future research to explore whether these amino acids correspond to common post-translational modifications or synthetic amino acids used in biocatalysis. Further exploration of these residues could provide insights into the model’s ability to design synthetic proteins with specialized, enhanced functions.

We show that the absorbing SELFIES model, in particular, excels in capturing complex chemical structures and generating novel, diverse sequences, indicating its promise for innovative protein design. However, challenges remain in enhancing the validity and structural reliability of the generated proteins.

Addressing these challenges is essential for translating computational designs into functional biological molecules. Future research should focus on refining the all-atom models to reduce the proportion of unusable sequences, addressing biases in amino acid distributions, and enhancing structural stability across varying sequence lengths. By overcoming these limitations, we can advance the design of all-atom proteins through generative models.

## Abbreviations

BFS: Breadth-first search
BLAST: Basic local alignment search tool
DDPM: Denoising diffusion probabilistic model
D3PM: Discrete denoising diffusion probabilistic model
D3FG: Functional-group-based diffusion model
DFM: Discrete flow model
GPT: Generative pre-trained transformer
GPU: Graphics processing unit
KL: Kullback–Leibler
mRNA: messenger Ribonucleic acid
NOS: Diffusion optimized sampling
pLDDT: predicted Local distance difference test
PTM: Post-translational modification
SELFIES: Self-referencing embedded string
SMILES: Simplified molecular-input line-entry system

## Declarations

## Acknowledgements

The authors would like to thank Mehul Bhuradia, Gabriel Vogel, and Lorenzo di Fruscia from the Intelligent Molecular Systems group at TU Delft for their insightful discussions, technical assistance, and constructive feedback throughout this work.

## Availability of data and materials

Data and code for reproducing this work are available at https://github.com/ Intelligent-molecular-systems/All-Atom-Protein-Sequence-Generation.

## Authors’ contributions

AVM and JMW conceived the idea; GJA curated the data, designed the all-atom analyses, implemented the models; AVM and GJA ran the experiments and wrote the manuscript; JMW, AVM, and MJTR supervised the project and reviewed the manuscript.

## Supplementary Material

### A Background

Here, we present the technical details of various design choices for our research. We first examine the all-atom molecular representations. Next, we discuss the general continuous diffusion model DDPM. Lastly, we present the ByteNet architecture used for the generative diffusion process.

#### A.1 All-Atom Molecular Representations

Proteins, being chains of bonded amino acids, are intuitively represented by their amino acid sequences. However, they can also be described by their detailed molecular structures. Representing a molecule as a linear string is challenging due to non-linear features like branches and rings. To address this, various techniques [1] have been developed, including SMILES [2], InChI [3], and deep-learning approaches such as DeepSMILES [4] and SELFIES [5].

Simplified molecular-input line-entry system (SMILES) strings [2] have been a prominent method for representing molecular graphs in computational chemistry since 1988. In SMILES, molecules are defined as sequences of atoms represented by letters, with branches denoted by parentheses and ring closures indicated by matching numbers. While SMILES grammar allows for the description of complex structures and properties like stereochemistry and chirality, it is not inherently robust; generative models can produce invalid strings that do not correspond to valid molecular graphs. To tackle this, self-referencing embedded strings (SELFIES) [5] offer a 100% robust molecular string representation, meaning that any combination of tokens corresponds to a chemically valid molecule. This robustness is achieved because SELFIES are designed to prevent the generation of syntactically and semantically invalid molecules by construction. This property is crucial in generative tasks where producing invalid sequences is undesirable.

In SELFIES, overloading is used to encode chemical structures in a way that eliminates common syntactic errors found in SMILES, such as unbalanced branch parentheses or incorrect ring identifiers. Overloading, in this context, means that certain tokens serve multiple purposes depending on their position and context in the sequence. For example, special tokens like [Branch1] or [Ring1] initiate branches or rings, and rather than requiring explicit end symbols, the subsequent tokens determine the length and connectivity of these features. This approach simplifies the representation and ensures structural validity throughout the sequence.

Moreover, the SELFIES grammar dynamically tracks the number of available bonds to prevent the generation of semantically incorrect molecules. If a sequence exhausts the available bonds, the grammar omits further tokens, ensuring the molecule remains chemically valid.

#### A.2 ByteNet Architecture

ByteNet [6] is a convolutional neural network (CNN) architecture designed for sequence-to-sequence tasks, such as machine translation. The architecture utilizes an encoder-decoder structure, where dilated convolutions are applied in the latent space, allowing the model to capture long-range dependencies within the sequence. Each sequence passes through multiple ByteNet blocks, where dilation functions act as a context window. A context window is the receptive field within which the model can “see” and process surrounding tokens in the sequence. It defines the number of tokens the model considers at a given position, helping it capture dependencies across various ranges without requiring recurrent processing. In each block, the dilation factor denoted as *k* increases exponentially for each subsequent layer, following the relation: *k* = 2^(*n*^ ^mod^ *^p^*^)^, where *n* is the layer index, and *p* = ⌊log_2_ *r*⌋ + 1, with *r* being the maximum dilation factor at the last block. This exponential growth in dilation allows ByteNet to efficiently cover long contexts in the sequence without increasing the number of layers, which improves the model’s efficiency and effectiveness for tasks involving long sequences.

The operations within each ByteNet block are shown in Fig. S1. The layers include normalization (LayerNorm) and activation functions (GeLU), followed by 1 × 1 convolutions and dilated convolutions with varying dilation factors. These operations ensure that the network captures both local and long-range features, while the residual connections enable efficient training by allowing information to bypass certain layers, facilitating gradient flow and avoiding vanishing gradients.

**Fig. S1:**
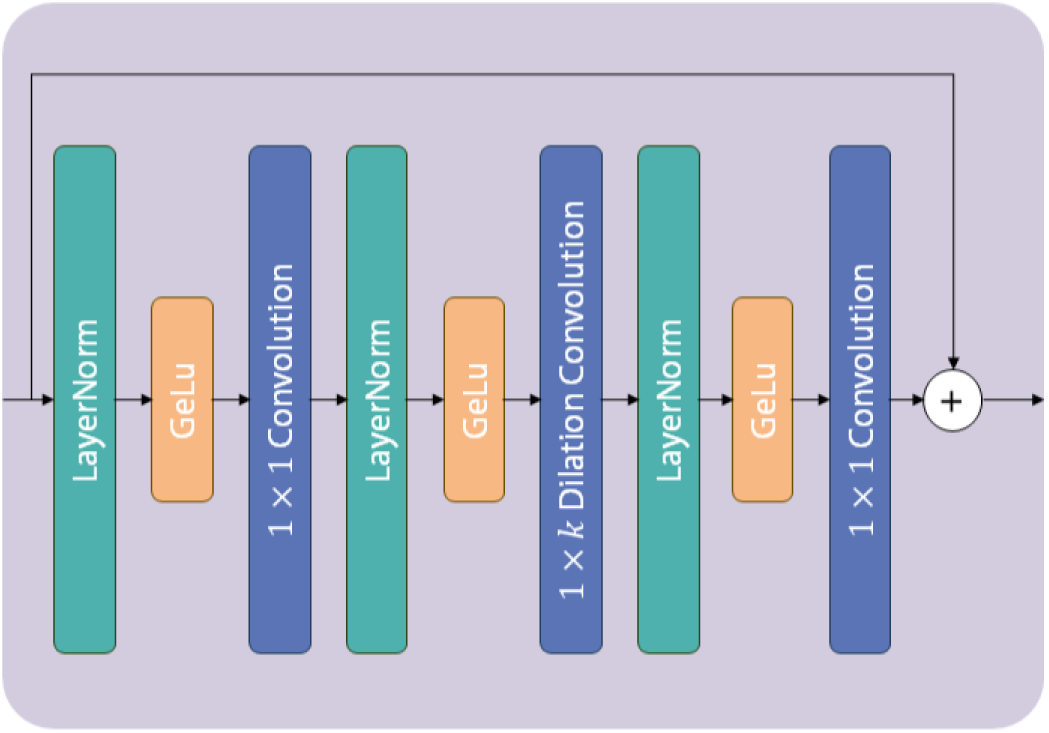
Schematic of a ByteNet block. Each block includes multiple operations: layer normalization, GeLU activations, 1×1 convolutions, and a key 1×*k* dilated convolution layer. The dilation factor, *k*, increases exponentially across layers, allowing the network to capture long-range dependencies. Residual connections are incorporated to aid in model training and gradient flow. (Adapted from [7])

ByteNet stands out for its ability to leverage parallel computation across sequences due to its fully convolutional design, making it highly efficient, especially when handling long input sequences. Unlike transformers, which experience quadratic scaling with sequence length and can become computationally intensive. Notably, studies have shown that ByteNet achieves comparable performance to transformers in tasks such as masked protein sequence modeling [7].

#### A.3 Denoising Diffusion Probabilistic Models (DDPMs)

Three sub-types of diffusion processes exist: denoising diffusion probabilistic models (DDPMs) [8], score-based generative models (SGMs) [9], and stochastic differential equations (SDEs) [10]. Diffusion processes aim to learn data distributions by gradually adding noise to the input and then learning how to reverse this process. While all diffusion models follow this general idea, they differ in how they add and remove noise, and in the architectures they use to reverse the noise. By learning how to effectively denoise corrupted inputs, these models can generate new, realistic data from the learned distributions.

In DDPMs, both the forward and backward processes are defined as Markov chains—a sequence of events where the state of the previous event dictates the probability of the next. A schematic overview of the whole DDPM process is given in Fig. S2.

In the DDPM framework, the forward diffusion process iteratively transforms the original distribution over a specified number of steps, denoted as *T*. This transformation gradually introduces noise, ultimately converging toward a simpler prior distribution, often a standard Gaussian distribution. The amount of noise added at each step is controlled by a predefined noise schedule, denoted as *β_t_*.

**Fig. S2:**
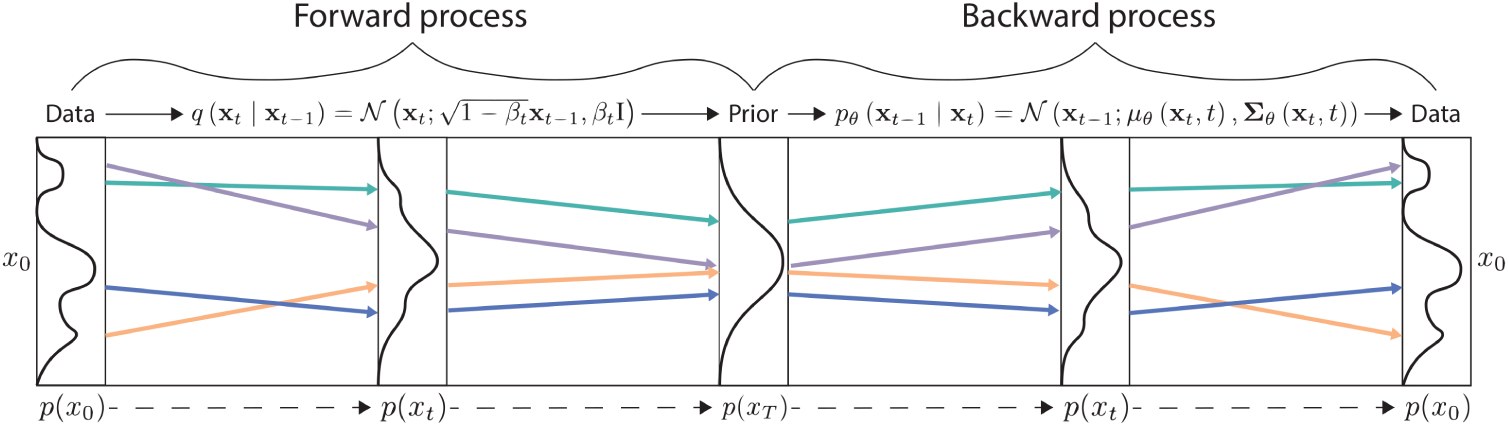
Schematic of a Denoising Diffusion Probabilistic Model (DDPM) using a continuous Gaussian noising process. This figure illustrates the forward process, which progressively transforms the original complex data into noise, and the backward process, which reverses the noise to generate new data samples. This enables us to generate novel data by learning the underlying distribution of the training data. (Adapted from [10])

Formally, the forward process is defined by the probability *q*(***x****_t_* | ***x****_t_*_−1_), where ***x****_t_* signifies the original input with noise corresponding to timestep *t*. When a DDPM is used with a continuous Gaussian noise process, its forward process is given as:

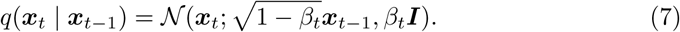

The backward diffusion process uses a neural network architecture with parameters *θ* that learns to predict the noise added in a forward step. This backward process reconstructs the original input based on the predicted noise at each timestep. The backward process with a Gaussian noise process is given as:

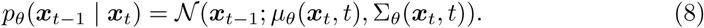

To optimize the generative model *p_θ_*(***x***_0_) and fit it to the data distribution *q*(***x***_0_), we minimize the variational upper bound on the negative log-likelihood:

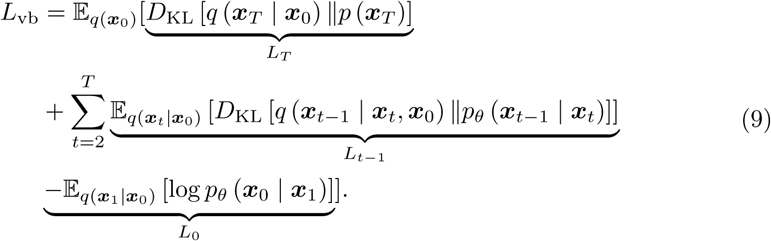

This equation represents the sum of Kullback–Leibler (KL) divergences between the forward and backward processes at each timestep. The KL divergence measures the statistical distance between a reference and a second probability distribution. The term *L_T_* represents the divergence at the final timestep of the process, while *L*_0_ is the reconstruction loss for the original data sample. The intermediate terms *L_t_*_−1_ account for the reconstruction terms between adjacent noising timesteps.

Lastly, selecting an appropriate prior distribution is crucial. The prior distribution must allow for a tractable forward posterior process *q*(***x****_t_*_−1_ | ***x****_t_,* ***x***_0_) to calculate the KL-divergence loss. Additionally, it must allow efficient computation of ***x****_t_* from ***x***_0_ using *q*(***x****_t_* | ***x***_0_) for any time *t*. These criteria are met when working with a standard Gaussian noise process.

### B Dataset Analysis

**Fig. S3:**
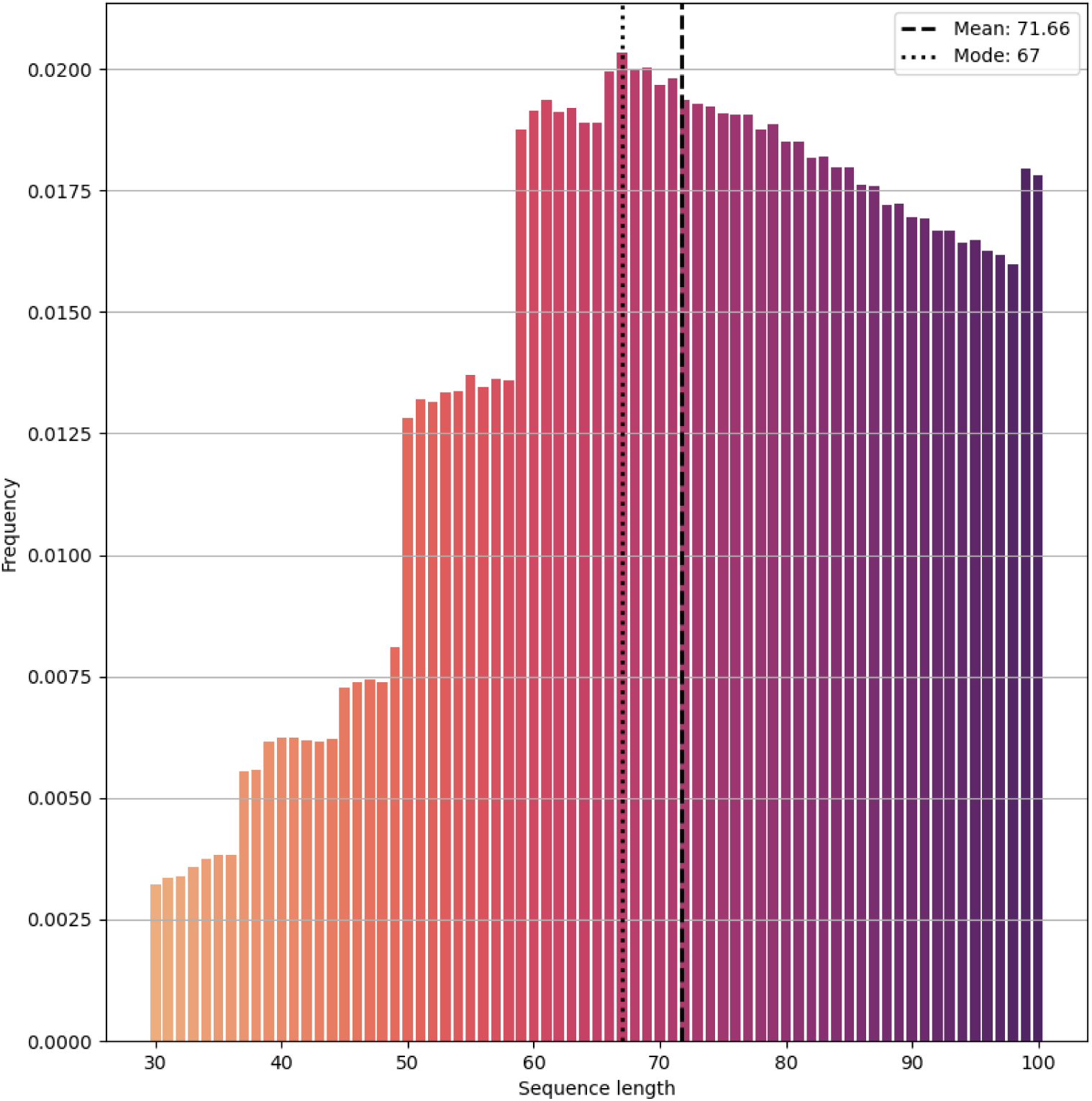
Distribution of protein sequence lengths in the amino acid representation, ranging from lengths 30 to 100. Both the mean and mode are approximately at length 70, indicating that most proteins in the dataset are around this length. There are relatively few proteins with lengths between 30 and 50. After peaking at length 70, the frequency of sequence lengths declines steadily from 70 to 98. Notably, there is a spike in frequency at lengths 99 and 100.

**Fig. S4:**
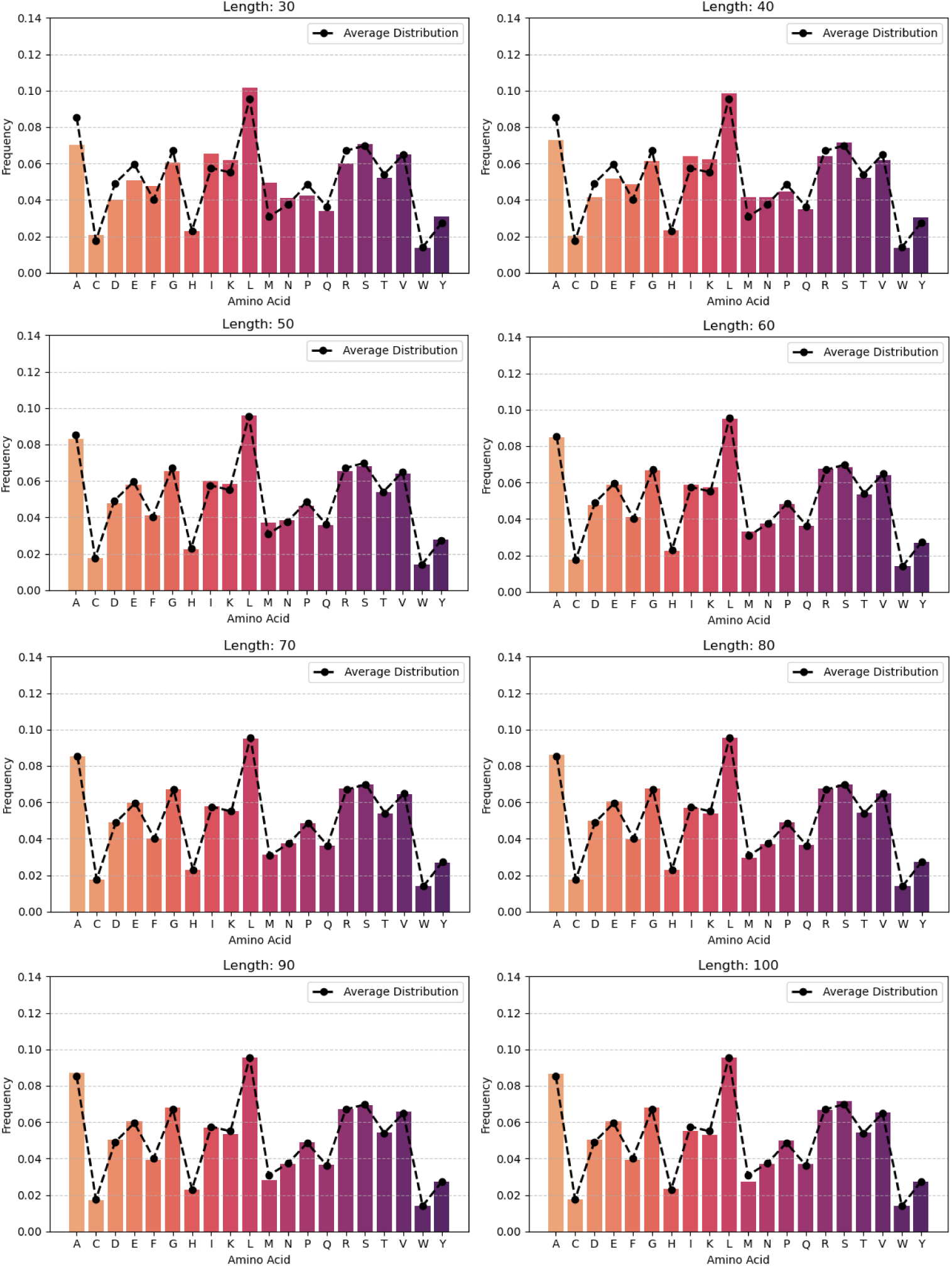
Amino acid token distributions for protein sequences grouped by length (30–40, 40–50, …, 90–100). The dotted line represents the average amino acid distribution for the full dataset. Sequences from 50–100 closely match the average distribution, reflecting their dominance in the dataset. Shorter sequences (30–50) show slight deviations but still reasonably approximate the overall dataset’s amino acid composition.

**Fig. S5:**
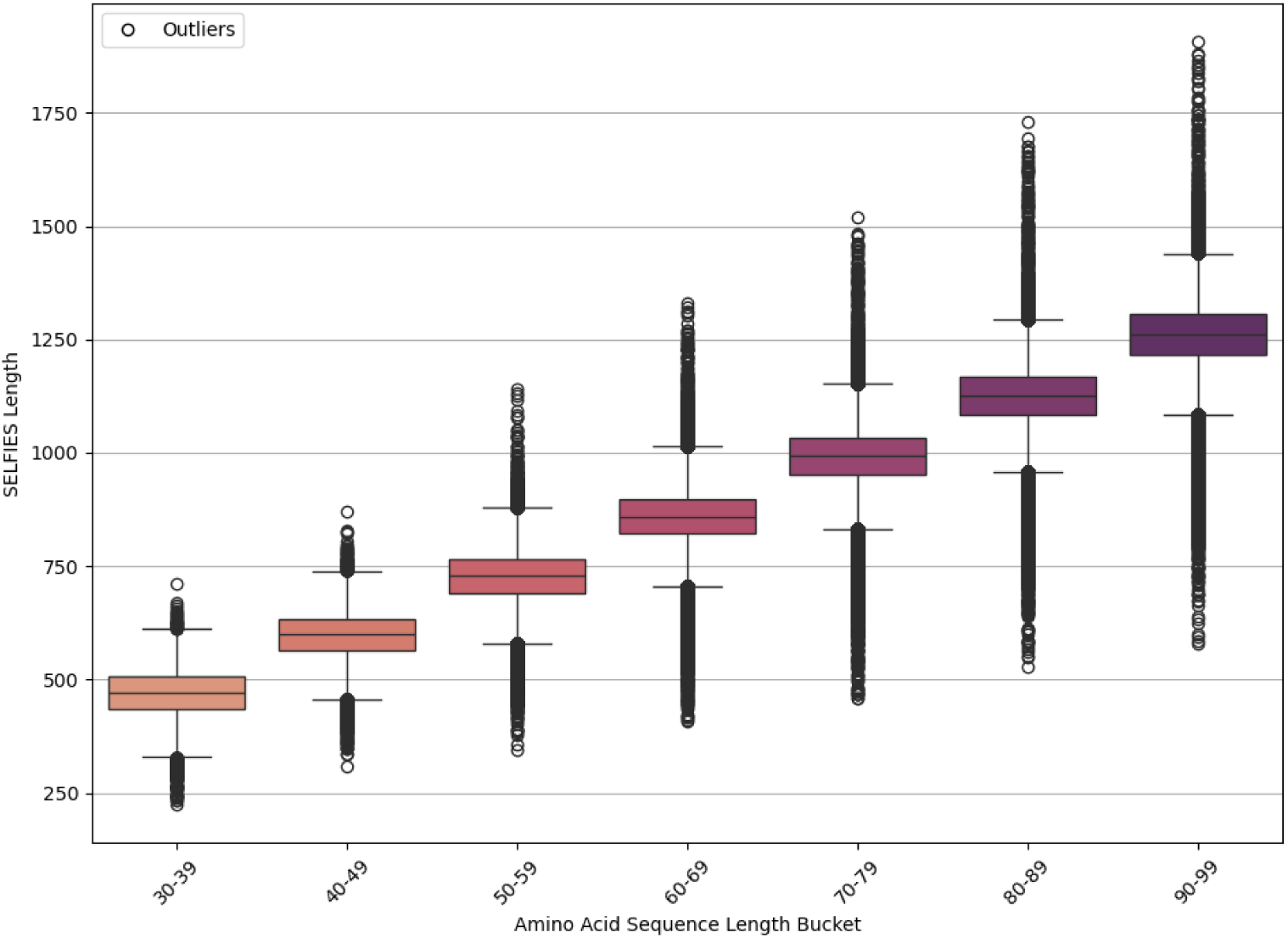
Distribution of SELFIES lengths by amino acid sequence length buckets. The box plots show that as amino acid sequence length increases, the average SELFIES length scales up approximately linearly. While the spread (interquartile range) increases gradually with longer sequences, indicating a relatively stable distribution, the number of outliers grows more noticeably. This suggests that while longer amino acid sequences generally correspond to longer SELFIES representations, there is an increasing degree of variability at these lengths.

### C SELFIES Sequences Denoising Process

**Table S1:**
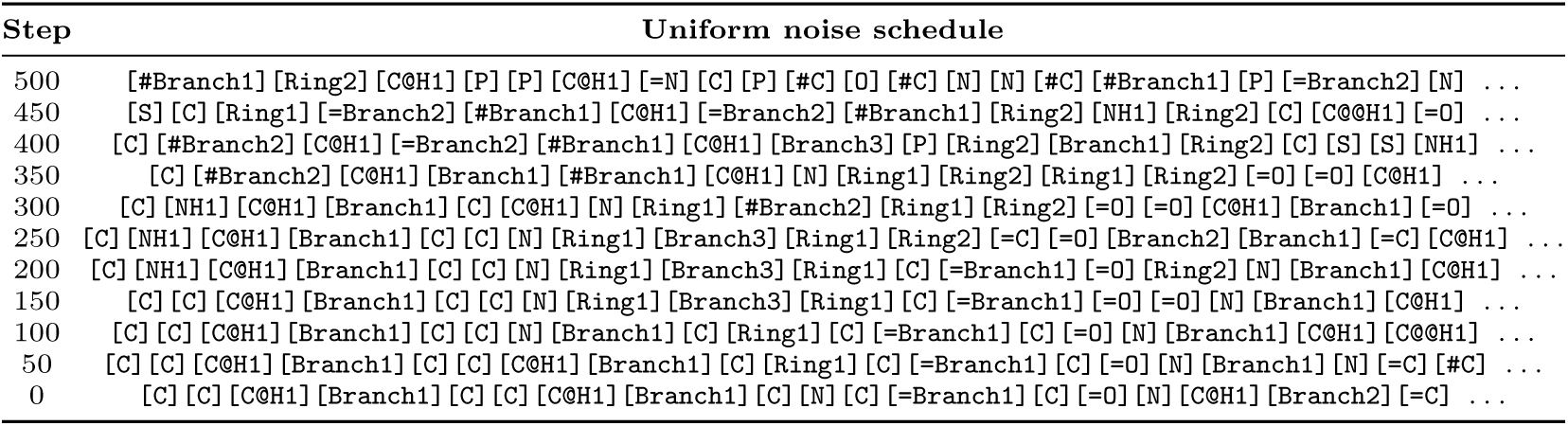
Sequence generation progression for the all-atom SELFIES representation using the uniform noise schedule.

**Table S2:**
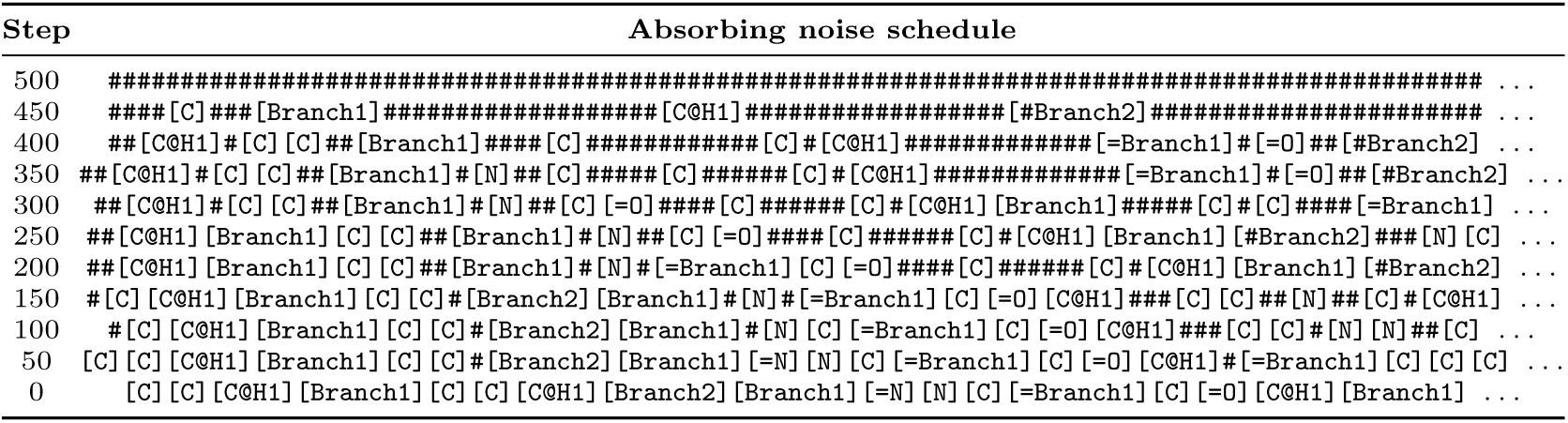
Sequence generation progression for the all-atom SELFIES representation using the absorbing noise schedule.

### D Extended All-Atom Results

**Table S3:** Unused SELFIES tokens and protein categorization for the 1000 unfiltered all-atom SELFIES sequences generated by the uniform and absorbing noise models. The results are shown for all SELFIES lengths (number of tokens) and grouped into seven evenly spaced length ranges.

|  | All<br>lengths | 225-465 | 465-705 | 705-945 | 945-1185 | 1185-1425 | 1425-1665 | 1665-1907 |
| --- | --- | --- | --- | --- | --- | --- | --- | --- |
| **Uniform model** |  |  |  |  |  |  |  |  |
| % Unused SELFIES tokens (↓) | 55.1% | 23.2% | 35.5% | 44.1% | 62.5% | 68.4% | 73.3% | 76.9% |
| N. Proteins with peptide bond (↑) | 980 | 128 | 152 | 139 | 131 | 134 | 150 | 146 |
| N. Proteins with continuous backbone (↑) | 52 | 28 | 12 | 7 | 3 | 1 | 1 | 0 |
| N. Non-canonical proteins (↑) | 44 | 24 | 10 | 6 | 3 | 0 | 1 | 0 |
| N. Canonical proteins (↑) | 4 | 3 | 1 | 0 | 0 | 0 | 0 | 0 |
| Non-canonical seq. length (avg.±std. ↑) | 24.1±13.3 | 21.8 | 21.9 | 39.0 | 25.0 | — | 9.0 | — |
| Canonical seq. length (avg.±std. ↑) | 30.5±7.2 | 27.0 | 41.0 | — | — | — | — | — |
| **Absorbing model** |  |  |  |  |  |  |  |  |
| % Unused SELFIES tokens (↓) | 18.9% | 8.0% | 10.9% | 13.4% | 20.0% | 21.8% | 25.4% | 31.8% |
| N. Proteins with peptide bond (↑) | 995 | 142 | 139 | 135 | 157 | 132 | 143 | 147 |
| N. Proteins with continuous backbone (↑) | 239 | 72 | 60 | 47 | 28 | 18 | 7 | 7 |
| N. Non-canonical proteins (↑) | 150 | 30 | 31 | 35 | 26 | 17 | 7 | 4 |
| N. Canonical proteins (↑) | 77 | 41 | 27 | 7 | 2 | 0 | 0 | 0 |
| Non-canonical seq. length (avg.±std. ↑) | 60.9±30.7 | 26.4 | 42.0 | 59.4 | 81.2 | 90.2 | 109.4 | 138.0 |
| Canonical seq. length (avg.±std. ↑) | 37.3±15.3 | 25.5 | 45.4 | 61.9 | 82.5 | — | — | — |

### E Distribution of Amino Acid Sequence Lengths

**Fig. S6:**
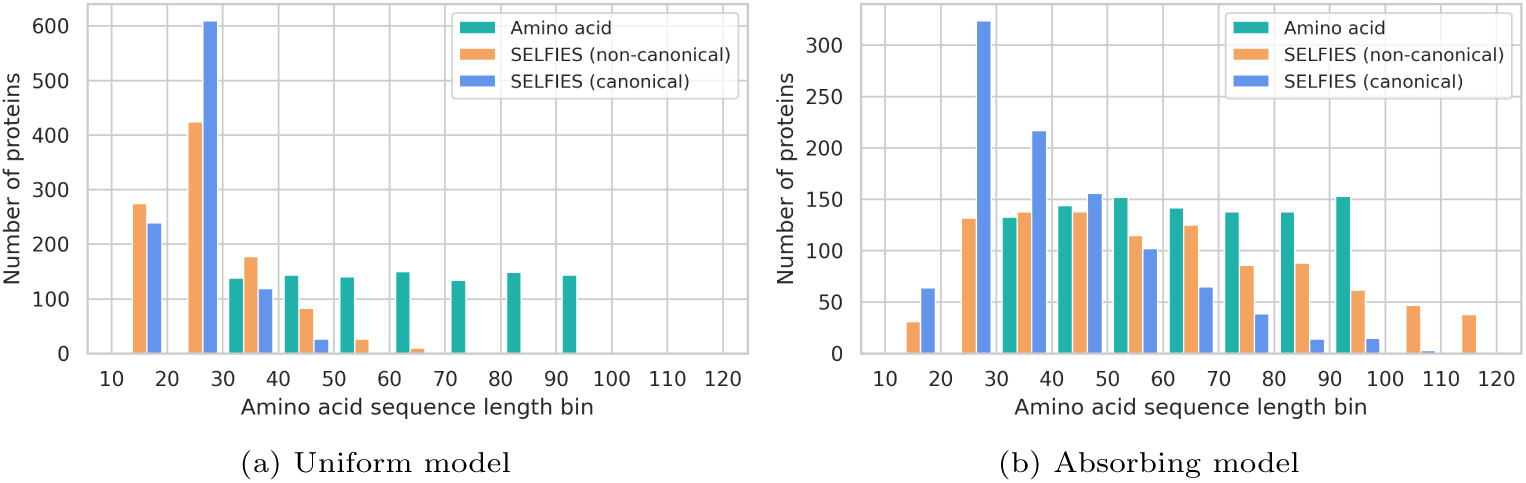
Distribution of amino acid sequence lengths for the 1000 valid proteins in each of the amino acid, SELFIES non-canonical, and SELFIES canonical sets, generated by the **(a)** uniform and **(b)** absorbing noise models. Protein counts are shown in evenly spaced bins spanning lengths from 10 to 120 amino acids.

### F Extended BLAST Results

Table S4 shows the BLAST results for 1000 amino acid sequences generated using both uniform and absorbing noise processes, evaluated across different e-value thresholds. A match with an e-value lower than 10^−5^ is considered significant. The table illustrates how varying the e-value threshold impacts the number of total matches and unique query IDs obtained for both novelty and diversity metrics. As the e-value threshold becomes less stringent (increasing from lower to higher values), both the total matches and unique query IDs increase for both noising processes. This analysis informs the selection of an e-value threshold of 0.05 for our final results, providing a balance between sensitivity and specificity in detecting significant sequence alignments.

**Table S4:**
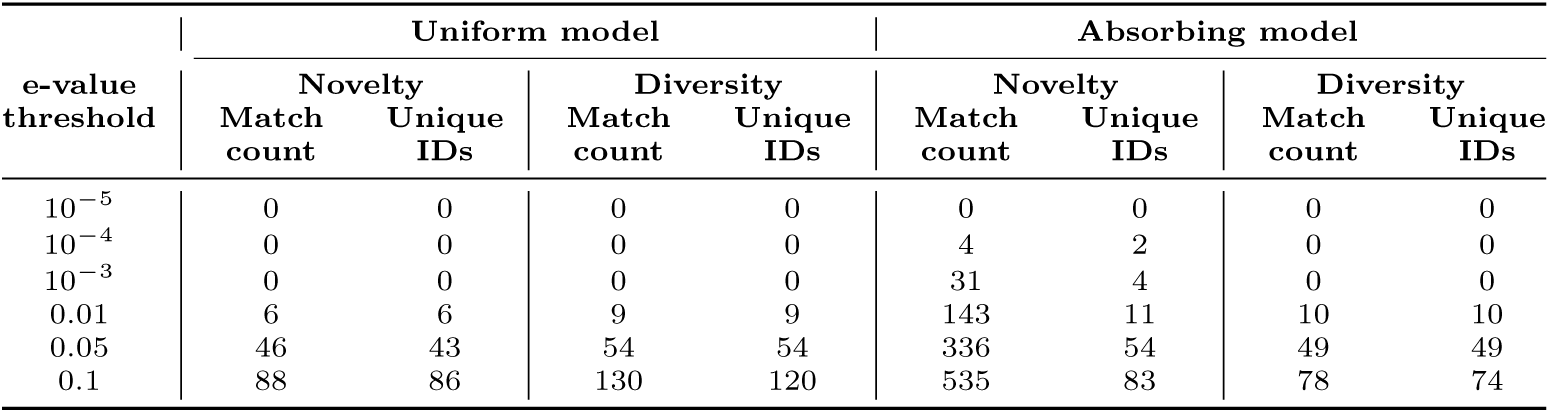
BLAST results for the 1000 amino acid sequences generated by the uniform and absorbing noise models, evaluated across different e-value thresholds.

| e-value threshold | Uniform model |  |  |  | Absorbing model |  |  |  |
| --- | --- | --- | --- | --- | --- | --- | --- | --- |
|  | Novelty |  | Diversity |  | Novelty |  | Diversity |  |
|  | Match count | Unique IDs | Match count | Unique IDs | Match count | Unique IDs | Match count | Unique IDs |
| $10^{-5}$ | 0 | 0 | 0 | 0 | 0 | 0 | 0 | 0 |
| $10^{-4}$ | 0 | 0 | 0 | 0 | 4 | 2 | 0 | 0 |
| $10^{-3}$ | 0 | 0 | 0 | 0 | 31 | 4 | 0 | 0 |
| 0.01 | 6 | 6 | 9 | 9 | 143 | 11 | 10 | 10 |
| 0.05 | 46 | 43 | 54 | 54 | 336 | 54 | 49 | 49 |
| 0.1 | 88 | 86 | 130 | 120 | 535 | 83 | 78 | 74 |

**Table S5:** BLAST results for novelty and diversity among the 1000 valid proteins in each of the amino acid, SELFIES non-canonical, and SELFIES canonical sets, generated by the uniform and absorbing noise models. Results are filtered to include matches with an e-value lower than 0.05. The average e-value ranges from 0.015 to 0.03. Lower is better for all the metrics.

| | N. Matches<br>(Unique matches) | % Sequences<br>with a match | Score<br>(avg. ± std.) | Query coverage<br>(avg. ± std.) | Identity<br>(avg. ± std.) |
| --- | --- | --- | --- | --- | --- |
| **Novelty** (e-value < 0.05) |  |  |  |  |  |
| **Uniform model** |  |  |  |  |  |
| Amino acid | 46 (43) | 4.3% | $80.0 \pm 2.8$ | $67.9 \pm 16.0$ | $16.9 \pm 2.6$ |
| SELFIES (non-canonical) | 1 (1) | 0.1% | $76.0 \pm 0.0$ | $86.8 \pm 0.0$ | $15.0 \pm 0.0$ |
| SELFIES (canonical) | 2 (2) | 0.2% | $76.5 \pm 1.5$ | $83.5 \pm 2.6$ | $14.5 \pm 1.5$ |
| **Absorbing model** |  |  |  |  |  |
| Amino acid | 336 (54) | 5.4% | $81.7 \pm 5.2$ | $66.5 \pm 15.3$ | $16.5 \pm 3.2$ |
| SELFIES (non-canonical) | 22 (22) | 2.2% | $81.5 \pm 4.6$ | $66.1 \pm 19.6$ | $18.9 \pm 4.2$ |
| SELFIES (canonical) | 15 (15) | 1.5% | $77.6 \pm 3.6$ | $80.1 \pm 13.8$ | $16.8 \pm 3.4$ |
| **Diversity** (e-value < 0.05) |  |  |  |  |  |
| **Uniform model** |  |  |  |  |  |
| Amino acid | 54 (54) | 5.4% | $51.0 \pm 3.2$ | $53.1 \pm 17.1$ | $11.3 \pm 2.4$ |
| SELFIES (non-canonical) | 3 (3) | 0.3% | $41.3 \pm 1.7$ | $55.4 \pm 4.6$ | $8.3 \pm 0.9$ |
| SELFIES (canonical) | 22 (22) | 2.2% | $40.9 \pm 2.2$ | $76.6 \pm 12.4$ | $8.5 \pm 1.4$ |
| **Absorbing model** |  |  |  |  |  |
| Amino acid | 49 (49) | 4.9% | $50.9 \pm 4.4$ | $52.2 \pm 19.2$ | $11.1 \pm 3.1$ |
| SELFIES (non-canonical) | 20 (19) | 1.9% | $50.4 \pm 2.9$ | $47.0 \pm 21.3$ | $12.1 \pm 1.8$ |
| SELFIES (canonical) | 36 (36) | 3.6% | $47.0 \pm 2.8$ | $60.1 \pm 21.8$ | $10.3 \pm 2.0$ |

### G Extended OmegaFold pLDDT Results

**Fig. S7:**
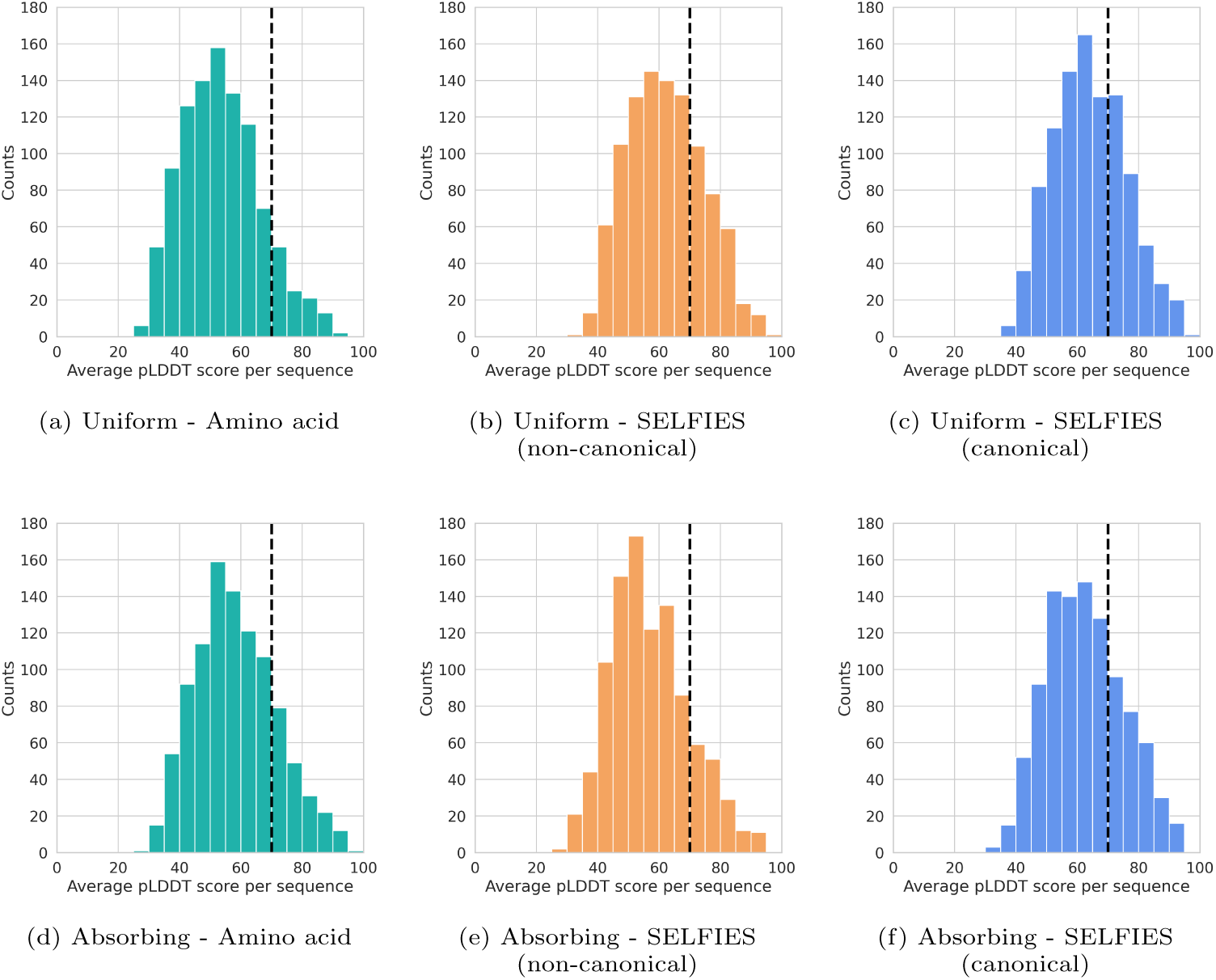
Distribution of OmegaFold average pLDDT scores for the 1000 valid proteins in each of the amino acid, SELFIES non-canonical, and SELFIES canonical sets, generated by the uniform and absorbing noise models. Dotted lines indicate the pLDDT threshold of 70, above which structure predictions are considered reliable.

**Fig. S8:**
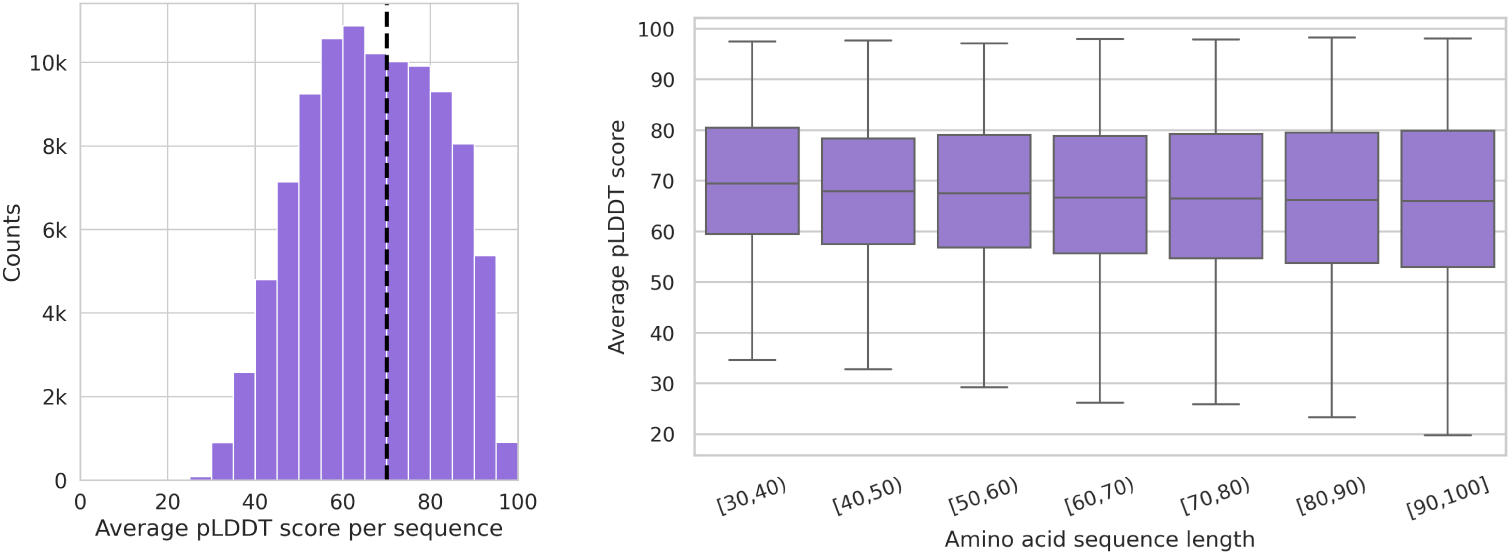
Distribution of OmegaFold average pLDDT scores for 100k training samples.

**Table S6:** OmegaFold average pLDDT confidence scores for the 1000 valid protein in each of the amino acid, SELFIES non-canonical, and SELFIES canonical sets, generated by the uniform and absorbing noise models. The results are shown for all lengths and grouped into nine length ranges. The first [10-30) and last [100-120) ranges handle outliers, while the rest are evenly spaced from 30 to 100, following the training set distribution. We also compare with the pLDDT score of 100k samples from the training set. Higher is better for the pLDDT score. Best results are highlighted in bold.

| | All<br>(avg.±std.) | [10-30) | [30-40) | [40-50) | [50-60) | [60-70) | [70-80) | [80-90) | [90-100) | [100-120) |
| --- | --- | --- | --- | --- | --- | --- | --- | --- | --- | --- |
| **Train (100k)** |  |  |  |  |  |  |  |  |  |  |
| N. Proteins | 100k | 0 | 4.2k | 6.9k | 13.7k | 19.5k | 19.2k | 17.9k | 18.5k | 0 |
| Avg. pLDDT | 66.9±15.3 | — | 69.5 | 67.9 | 67.7 | 67.0 | 66.7 | 66.2 | 66.0 | — |
| N. pLDDT > 70 | 43.6k | — | 2.1k | 3.1k | 6.1k | 8.4k | 8.2k | 7.7k | 7.9k | — |
| **Uniform model** |  |  |  |  |  |  |  |  |  |  |
| **Amino acid** |  |  |  |  |  |  |  |  |  |  |
| N. Proteins | 1000 | 0 | 138 | 144 | 141 | 150 | 134 | 149 | 144 | 0 |
| Avg. pLDDT | 53.7±12.7 | — | 64.0 | 60.3 | 56.4 | 53.6 | 50.0 | 46.6 | 45.0 | — |
| N. pLDDT > 70 | 110 | — | 39 | 27 | 20 | 13 | 5 | 5 | 1 | — |
| **SELFIES** |  |  |  |  |  |  |  |  |  |  |
| **(non-canonical)** |  |  |  |  |  |  |  |  |  |  |
| N. Proteins | 1000 | 700 | 178 | 83 | 26 | 10 | 3 | 0 | 0 | 0 |
| Avg. pLDDT | 62.3±12.3 | 64.7 | 58.3 | 54.2 | 54.0 | 53.6 | 51.9 | — | — | — |
| N. pLDDT > 70 | 272 | 228 | 31 | 9 | 3 | 1 | 0 | — | — | — |
| **SELFIES** |  |  |  |  |  |  |  |  |  |  |
| **(canonical)** |  |  |  |  |  |  |  |  |  |  |
| N. Proteins | 1000 | 850 | 119 | 26 | 3 | 1 | 0 | 0 | 1 | 0 |
| Avg. pLDDT | **64.2±11.9** | 65.3 | 59.3 | 54.7 | 55.6 | 56.2 | — | — | 46.9 | — |
| N. pLDDT > 70 | **321** | 296 | 21 | 3 | 1 | — | — | 0 | — | — |
| **Absorbing model** |  |  |  |  |  |  |  |  |  |  |
| **Amino acid** |  |  |  |  |  |  |  |  |  |  |
| N. Proteins | 1000 | 0 | 133 | 144 | 152 | 142 | 138 | 138 | 153 | 0 |
| Avg. pLDDT | 58.5±13.1 | — | **65.5** | **63.9** | **60.9** | **57.8** | **55.1** | 53.0 | **53.4** | — |
| N. pLDDT > 70 | 194 | — | 45 | 47 | 32 | 26 | 16 | 12 | 16 | — |
| **SELFIES** |  |  |  |  |  |  |  |  |  |  |
| **(non-canonical)** |  |  |  |  |  |  |  |  |  |  |
| N. Proteins | 1000 | 163 | 138 | 138 | 115 | 125 | 86 | 88 | 62 | 85 |
| Avg. pLDDT | 57.0±12.7 | 67.3 | 62.2 | 60.5 | 55.7 | 54.3 | 53.9 | 48.9 | 48.3 | 46.2 |
| N. pLDDT > 70 | 162 | 60 | 45 | 29 | 12 | 7 | 6 | 0 | 2 | 1 |
| **SELFIES** |  |  |  |  |  |  |  |  |  |  |
| **(canonical)** |  |  |  |  |  |  |  |  |  |  |
| N. Proteins | 1000 | 388 | 217 | 156 | 102 | 65 | 39 | 14 | 15 | 4 |
| Avg. pLDDT | 62.7±12.5 | **68.3** | 62.0 | 61.0 | 57.8 | 53.1 | **55.1** | **56.3** | 50.1 | **49.8** |
| N. pLDDT > 70 | 279 | 177 | 44 | 39 | 12 | 3 | 4 | 0 | 0 | 0 |

### H Model Hyperparameters

**Table S7:** Summary of hyperparameter configurations used in our D3PM implementation with the ByteNet architecture, including details on the dataset processing, optimizer settings, and learning rate scheduler. Each hyperparameter is listed alongside its value and source.

| Hyperparameter | Value | Source |
| --- | --- | --- |
| **ByteNet model** |  |  |
| Embedding dimension ($d_{\text{embed}}$) | 8 | EvoDiff |
| Model dimension ($d_{\text{model}}$) | 1024 | EvoDiff |
| Activation function | GeLU | EvoDiff |
| Slim | True | EvoDiff |
| Number of layers ($n_{\text{layers}}$) | 16 | EvoDiff |
| Kernel size | 5 | EvoDiff |
| Max dilation value ($r$) | 128 | EvoDiff |
| Diffusion timesteps | 500 | EvoDiff |
| Number of tokens amino acid ($n_{\text{tokens}}$) | 20 | This work |
| Number of tokens SELFIES ($n_{\text{tokens}}$) | 21 | This work |
| Loss reweighting uniform ($\lambda$) | 0 | D3PM |
| Loss reweighting absorbing ($\lambda$) | 0.1 | D3PM |
| Causal | False | EvoDiff |
| Dropout | 0.1 | EvoDiff |
| Tie weights | False | EvoDiff |
| Final norm | False | EvoDiff |
| **Dataset and batch sampling** |  |  |
| Dataset | UniRef50 | This work |
| Max tokens | 40000 | EvoDiff |
| Max batch size | 800 | EvoDiff |
| Bucket size | 1000 | EvoDiff |
| Max epoch | 500 | EvoDiff |
| **Optimizer and scheduler** |  |  |
| Optimizer | Adam | EvoDiff |
| Learning rate | $10^{-4}$ | EvoDiff |
| Weight decay | 0 | EvoDiff |
| Scheduler | LambdaLR | EvoDiff |
| Warm-up steps | 10000 | EvoDiff |

### I Training and Validation Loss

**Fig. S9:**
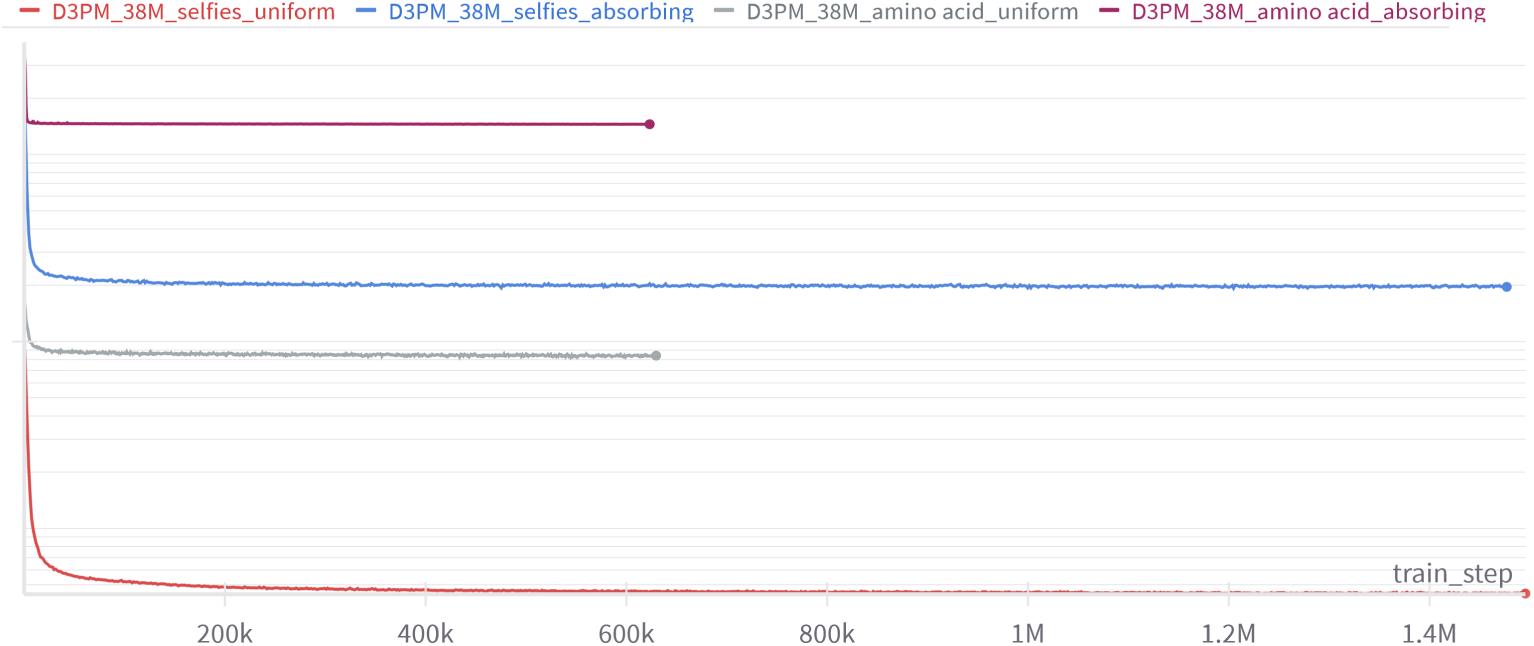
Training loss per step for the amino acid and SELFIES models under both uniform and absorbing noise schedules. All models have largely converged. (y-axis is in log scale).

**Fig. S10:**
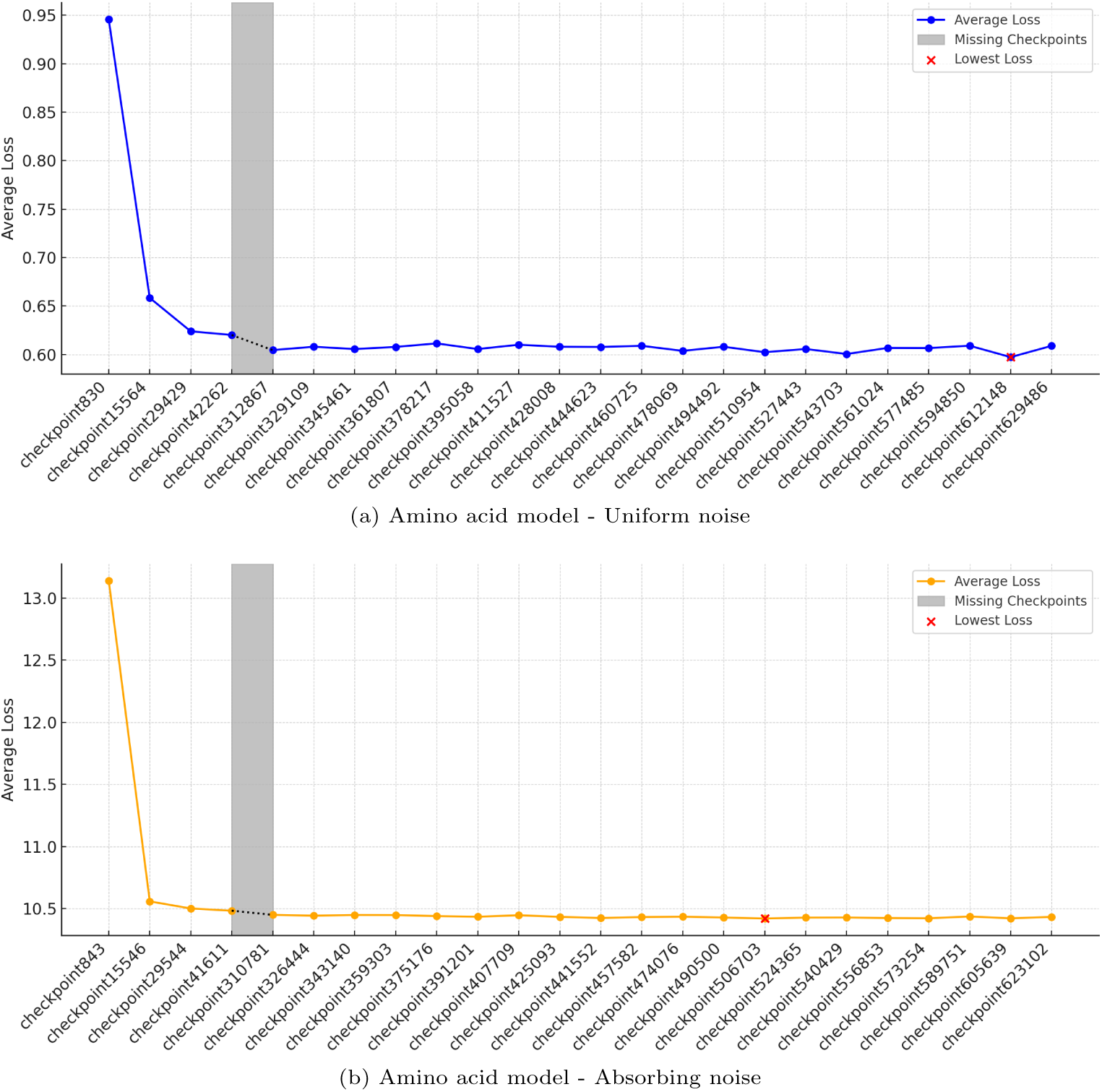
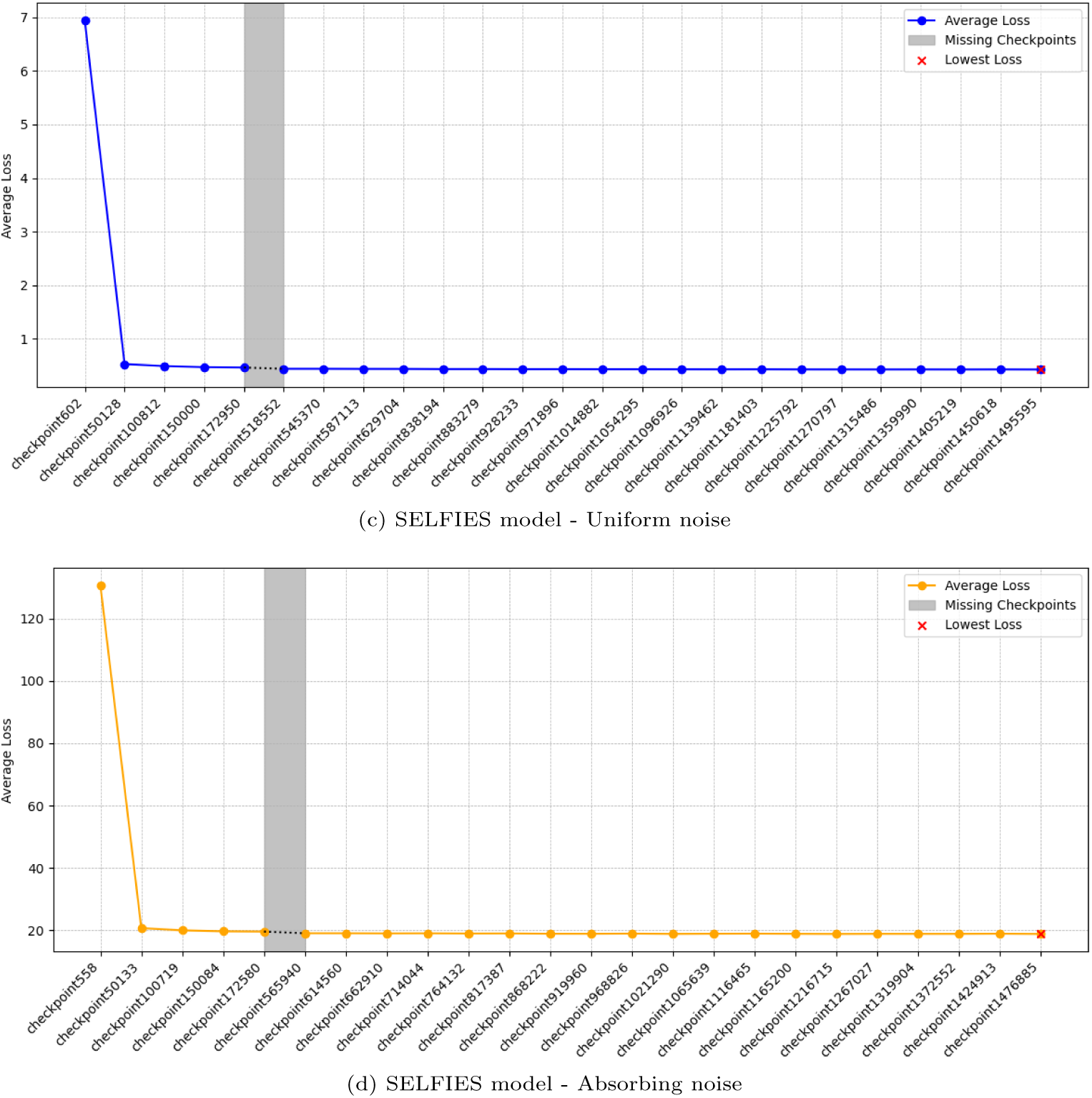
Average validation loss across different checkpoints for the amino acid and SELFIES models using both uniform and absorbing noise schedule. The lowest average validation loss checkpoint is highlighted, indicating the best-performing model. The trend shows that the models have largely converged, with minimal variation in loss across the later checkpoints.

